# Pulmonary fibroblast activation during *Aspergillus fumigatus* infection enhances lung defense via immunomodulation and tissue remodeling

**DOI:** 10.1101/2024.12.19.629499

**Authors:** José P. Guirao-Abad, Shannon M. Shearer, Jon Bowden, Daniel A. Kasprovic, Christina Grisham, Mustafa Ozdemir, Michael Tranter, Yunguan Wang, David S. Askew, Onur Kanisicak

## Abstract

*Aspergillus fumigatus* is the etiologic agent of invasive aspergillosis, a life-threatening fungal pneumonia that is initiated by the inhalation of conidia (spores) into the lung. If the conidia are not cleared, they secrete large quantities of hydrolytic enzymes and toxins as they grow, resulting in extensive damage to pulmonary tissue. Stromal fibroblasts are central responders to tissue damage in many organs, but their functional response to pulmonary injury caused by *A. fumigatus* has not been explored. In this study, we employed cell lineage tracing, targeted cell ablation, and single-cell RNA sequencing to monitor the dynamics of fibroblast behavior upon exposure to *A. fumigatus* in both immunocompetent and immunosuppressed hosts. The results demonstrate that a subset of pulmonary fibroblasts becomes activated in an immunocompetent host in response to a challenge with *A. fumigatus* conidia, acquiring a gene expression program reflecting the acquisition of new immunomodulatory properties as well as enhanced extracellular matrix (ECM)-secreting ability. Remarkably, through targeted ablation of fibroblasts that express the profibrotic activation marker periostin, we demonstrate that the progression of an invasive *A. fumigatus* infection in an immunosuppressed host is accelerated by the absence of periostin lineage cells and is accompanied by severe alveolar hemorrhage and angioinvasion. These findings uncover a novel protective role for fibroblasts in limiting the severity of *A. fumigatus*-induced pulmonary injury and emphasize the importance of the pulmonary stroma in host defense against this invasive fungal infection.

## INTRODUCTION

Environmental molds such as *Aspergillus fumigatus* commonly disseminate through the release of conidia (spores) into the air, inevitably exposing the lung to potential infectious threats^1^. Among these fungi, *A. fumigatus* stands out as one of the major invasive fungal infections (IFI) of global concern^2^. The outcome of the interaction between *A. fumigatus* and the lung hinges on the host’s immune status. While healthy immunocompetent lungs can successfully eradicate the inhaled conidia before they can initiate infection^3^, individuals with mild immunosuppression or those with pre-existing lung structural defects are vulnerable to pulmonary colonization by the fungus. These colonizing infections are generally not invasive, but the constitutive release of fungal hydrolytic enzymes and toxins triggers a cycle of repeated tissue injury and inflammation that may evolve into a long-term condition known as chronic pulmonary aspergillosis^4,5^. In situations of more severe immunosuppression that may arise in transplant patients, the conidia germinate into the invasive hyphal form of the fungus, resulting in a life-threatening infection known as invasive aspergillosis (IPA). IPA is characterized by the polarized growth of hyphae across anatomic boundaries, resulting in extensive tissue damage and dissemination to other tissues^4^. Infections caused by *A. fumigatus* are associated with a high rate of morbidity and mortality, even when treated, and the recent emergence of antifungal drug resistance^6^ underscores the need for more understanding of mechanisms involved in pulmonary protection.

The frontline of host defense against inhaled *A. fumigatus* spores involves the integrated activity of lung-resident macrophages and dendritic cells, along with recruited inflammatory cells^7–9^. However, little is known about the contribution of pulmonary stromal cells, defined here as support cells that lack hematopoietic, endothelial and epithelial markers, most of which are fibroblasts. Fibroblasts, which are best known for their role in providing structural support to a tissue through the synthesis and remodeling of the extracellular matrix (ECM), are increasingly recognized as a heterogeneous collection of matrix-secreting cells that can shift under pathologic conditions into distinct populations with diverse functionalities^10–12^. However, the dynamics of fibroblast behavior during the *Aspergillus*-host interaction is currently unexplored.

In this study, we genetically traced fibroblast activation using the previously identified activation marker periostin (Postn)^13–16^ and investigated the impact of these Postn-lineage (Postn^Lin^) fibroblasts on the pathogenesis of IPA. The findings reveal that pulmonary inoculation of immunocompetent mice with *A. fumigatus* conidia induces Postn expression. These Postn^Lin^ cells were identified as members of the stromal fibroblast population by both immunohistochemistry and bulk RNA-sequencing (RNA-seq) and were localized to anatomic sites expected for pulmonary fibroblasts. Although these Postn^Lin^ fibroblasts represented a subset of the total stromal cell population, their elimination in an inducible cell depletion model accelerated the progression of tissue damage and mortality during IPA. Singe-cell RNA sequencing (scRNA-seq) of lung stromal cell populations demonstrated that exposure to *A. fumigatus* induced the transient appearance of two novel clusters of cells characterized by the expression of distinct matrix-preserving genes, as well as pattern-recognition receptors and proinflammatory cytokines and chemokines with well-established roles in host defense against IFI’s. Together, these findings highlight a dual role for fibroblasts in pulmonary protection against *A. fumigatus*, involving both immunomodulation and ECM preservation.

## RESULTS

### Collagen deposition is localized at sites of inflammation induced by *A. fumigatus*

To assess the pulmonary response to *A. fumigatus*, we inoculated immunocompetent mice with a non-lethal dose of conidia (strain CEA10) via the oropharyngeal route (OP)^17^. As expected for immunocompetent animals^18^, colony-forming unit (CFU) analysis revealed sterilization of most conidia within 72 h (Fig. S1A). Histologic evaluation of lung tissue on day 1 post-inoculation revealed multiple inflammatory foci, which persisted for at least 7 days (Fig. 1A). However, the inflammatory foci were reduced in size on day 7 relative to day 3, suggesting the beginning of a resolution of the inflammatory response concomitant with fungal clearance. GMS-staining on day 1 post-inoculation demonstrated the presence of both conidia and small germlings within the inflammatory foci (Fig. 1B and S1C), which decreased by day 3 and only fungal debris was detected by days 5 and 7 (Fig. S1C). Trichrome staining revealed collagen fibers interspersed between the inflammatory cells on days 1, 3, 5, and 7 post-inoculation (Fig. 1C). Control sections from saline-treated mice are shown in Fig. S1B. We conclude that the accumulation of collagen within inflammatory foci is an initial response to a pulmonary challenge with *A. fumigatus* conidia in immunocompetent animals.

**Figure 1.**
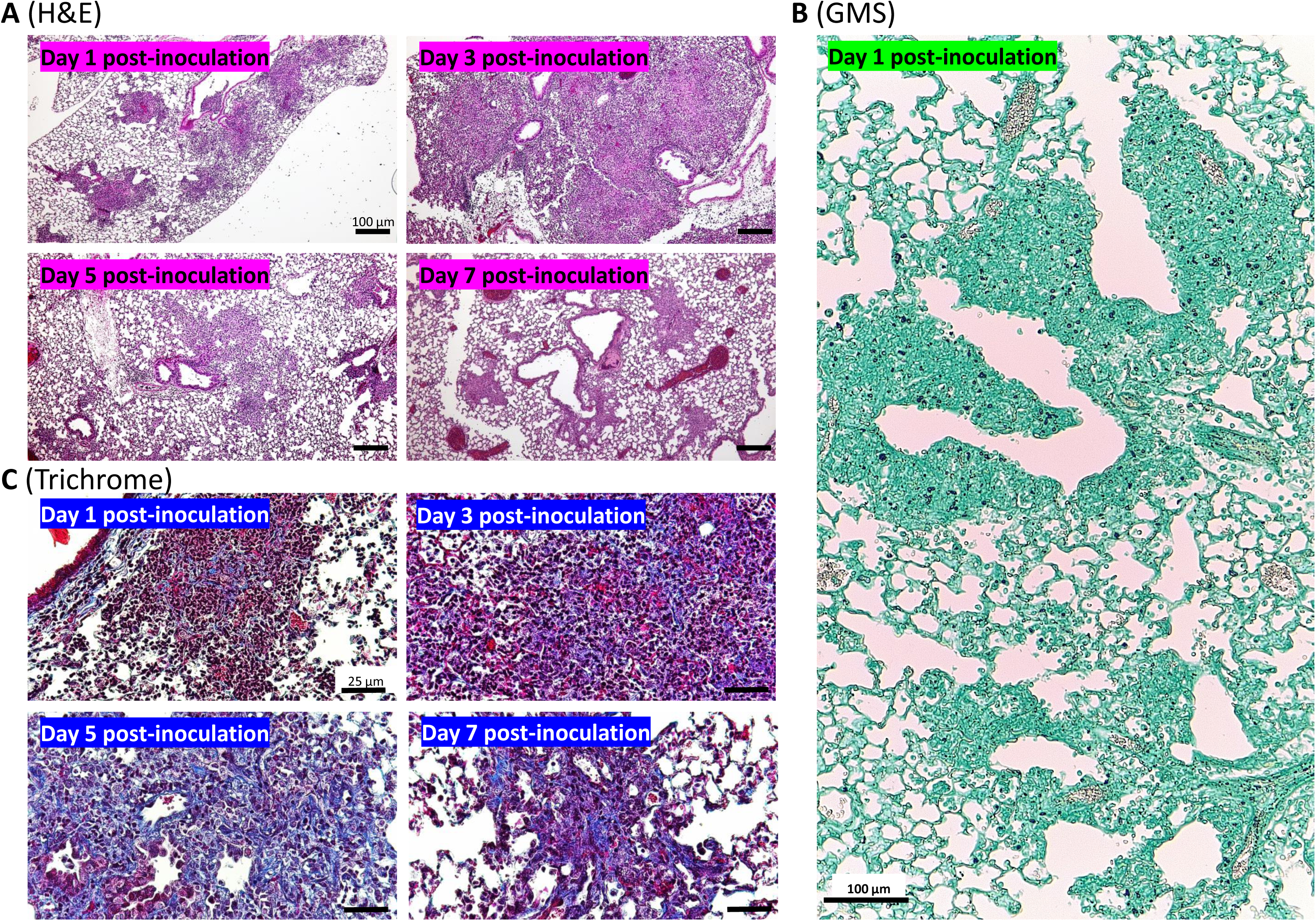
Collagen fiber deposition is an initial response to *A. fumigatus.* Representative images of paraffin-embedded lung sections from immunocompetent mice on days 1, 3, 5, and 7 post-inoculation with 3 x 10^7^ *A. fumigatus* conidia. Sections were stained with (A) hematoxylin and eosin (H&E), (B) Gomori’s methenamine silver (GMS) to highlight fungal elements (black), or (C) Masson’s trichrome to highlight collagen fibers (blue).

### Postn^Lin^ fibroblasts emerge in response to *A. fumigatus*

Detecting increased collagen deposition at sites of the fungus-host interaction implies that *A. fumigatus* stimulates fibroblasts to enhance ECM secretion. Changes in fibroblast activity in the context of tissue injury has been reported in many tissues and is associated with the induction of a gene expression program that modifies the collagen-producing activity of fibroblasts to maintain tissue integrity^19^. The periostin (Postn) gene, which encodes a matricellular protein involved in ECM organization, tissue remodeling, and the regulation of inflammatory cell migration^13–16^, is a well-established fibroblast activation marker. Most resident fibroblasts in adult tissues lack Postn expression, but they will activate the gene in response to injury^16^. In this study, we used a genetic cell lineage tracing approach to identify fibroblasts in which the Postn promoter has been induced in response to *A. fumigatus*.

To monitor Postn promoter activation, one Postn locus was replaced with a tamoxifen (TAM)-inducible Cre recombinase (Postn-MerCreMer) via targeted insertion, and crossed with a strain that permanently expresses the tdTomato fluorescent reporter gene in response to Cre activity (Fig. 2A). These Postn^Lin^-tracing mice were given an initial oral gavage of TAM on day -1 and then inoculated OP with a non-lethal dose of 3 x 10^7^ conidia in saline on day 0 (Fig. 2B). Only sporadic tdTomato+ fluorescence was observed in lung tissue from Postn^Lin^ mice treated with either TAM alone or TAM plus an OP inoculation of saline (Fig. 2C, top two panels and Fig. S2A, top 3 panels; arrows highlight tdTomato+ cells), representing background levels. No sporadic tdTomato expression was detected in the Postn^Lin^ mice in the absence of TAM treatment (Fig. S2A, lower left panel), indicating tight control of the reporter gene without Cre recombination. By contrast, robust tdTomato expression within a subset of fibroblasts was apparent in lungs inoculated with conidia from the CEA10 strain of *A. fumigatus* (Fig. 2C, lower left panel and Fig. S2A, lower middle panel). Interestingly, heat-inactivation of the CEA10 conidia induced a similar level of activation to viable conidia, indicating that metabolic activity of the fungus is not required for this response (Fig. S2A, lower right panel). Comparable levels of expression were induced by two other *A. fumigatus* clinical isolates (Af293 and H237), demonstrating that the response is not *A. fumigatus* strain-specific (Fig S2B). No expression was seen in mice inoculated with *A. fumigatus* without TAM treatment, demonstrating tight control of the reporter even with a strong inducer of fibroblast activation (Fig 2C, lower right panel). TdTomato-expressing activated fibroblasts were first detectable histologically on day 3 post-inoculation, with the number of labeled cells peaking on day 7 (Fig. 2D, quantified in 2E), reaching levels that were 100-fold higher, on average, than the sporadic activation observed following OP saline inoculation (Fig. 2F). The optimum dose range required to elicit a detectable activation response was 2-4 x 10^7^ conidia administered as a single OP inoculation (Fig. S3). We conclude that the *A. fumigatus*-host interaction in the immunocompetent lung triggers the emergence of a population of fibroblasts characterized by expression of the Postn gene.

**Figure 2.**
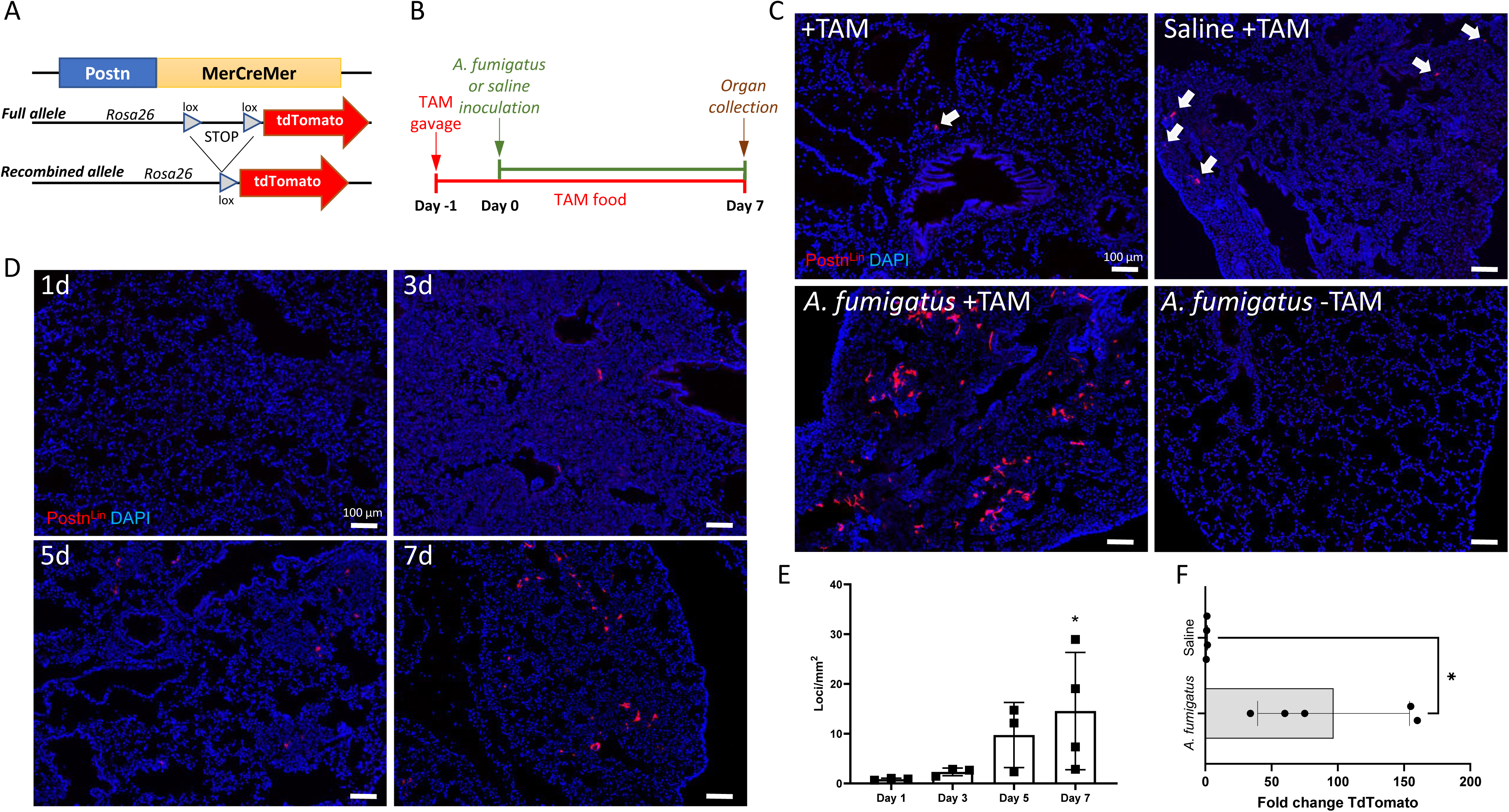
Postn^Lin^ cells are induced by *A. fumigatus* in immunocompetent mice. (A) Schematic representation of the TAM-inducible Cre recombinase system. Cre recombinase, flanked by a tamoxifen-inducible modified estrogen receptor (MerCreMer), was inserted into the Postn locus, enabling temporal control over Cre activity in cells that activate the Postn promoter. In the presence of TAM, Cre translocates to the nucleus and catalyzes recombination between loxP sites flanking a transcriptional stop upstream of a tdTomato reporter gene inserted into the rosa26 locus. Cre activity results in constitutive expression of tdTomato in Cre-expressing cells and their progeny, thereby fluorescently marking Postn^Lin^ fibroblasts. (B) Schematic representation of the experimental approach used to study fibroblast activation in Postn^Lin^ immunocompetent mice after OP inoculation with 3 x 10^7^ *A. fumigatus* conidia. TAM was administered via gavage one day before the fungal challenge and maintained through a TAM-infused chewing diet for the duration of the experiment. (C) Representative fluorescent photomicrographs showing the induction of tdTomato-positive cells following exposure to the specified treatments. Arrows indicate sporadic Postn^Lin^ cells in samples not treated with *A. fumigatus*. (D & E) Time course illustrating the appearance of tdTomato+ cells on days 1, 3, 5, and 7 post-inoculation with *A. fumigatus* (* P < 0.05; one-way ANOVA on ranks with Tukey’s post hoc test). (F) Flow cytometric analysis of tdTomato expression following inoculation with 3 x 10^7^ conidia or mock infection with saline (* P < 0.05; unpaired two-tailed t-test).

### Postn^Lin^ fibroblasts induced by *A. fumigatus* are predominantly alveolar

Characterization of fibroblast heterogeneity in the lung by scRNA-seq has revealed unique molecular profiles for populations of fibroblasts that reside in distinct anatomic sites: directly beneath the epithelial lining of large and small conducting airways (peribronchial fibroblasts), in the connective tissue surrounding bronchovascular bundles (adventitial fibroblasts), or within the alveolar interstitial space between epithelial walls (alveolar fibroblasts)^19^. To determine where Postn^Lin^ fibroblasts localize, we quantified the number of tdTomato+ cells at each site by immunohistochemistry, confirming localization by costaining with antibodies against epithelial (EpCAM), endothelial (CD31), and smooth muscle (Myh11) markers. The majority of the Postn^Lin^ cells arising 7 days post-inoculation with *A. fumigatus* conidia were found within the alveolar interstitium (Fig. 3B, arrows). A smaller proportion was sandwiched between the bronchiolar epithelium and the underlying smooth muscle (Fig. 3C, enlarged in Fig. 3D), and the fewest Postn^Lin^ cells showed an adventitial localization surrounding perivascular smooth muscle (Fig. 3E, enlarged in Fig. 3F). Quantification of TdTomato+ cells at the three major anatomic sites is shown in Fig. 3A.

**Figure 3.**
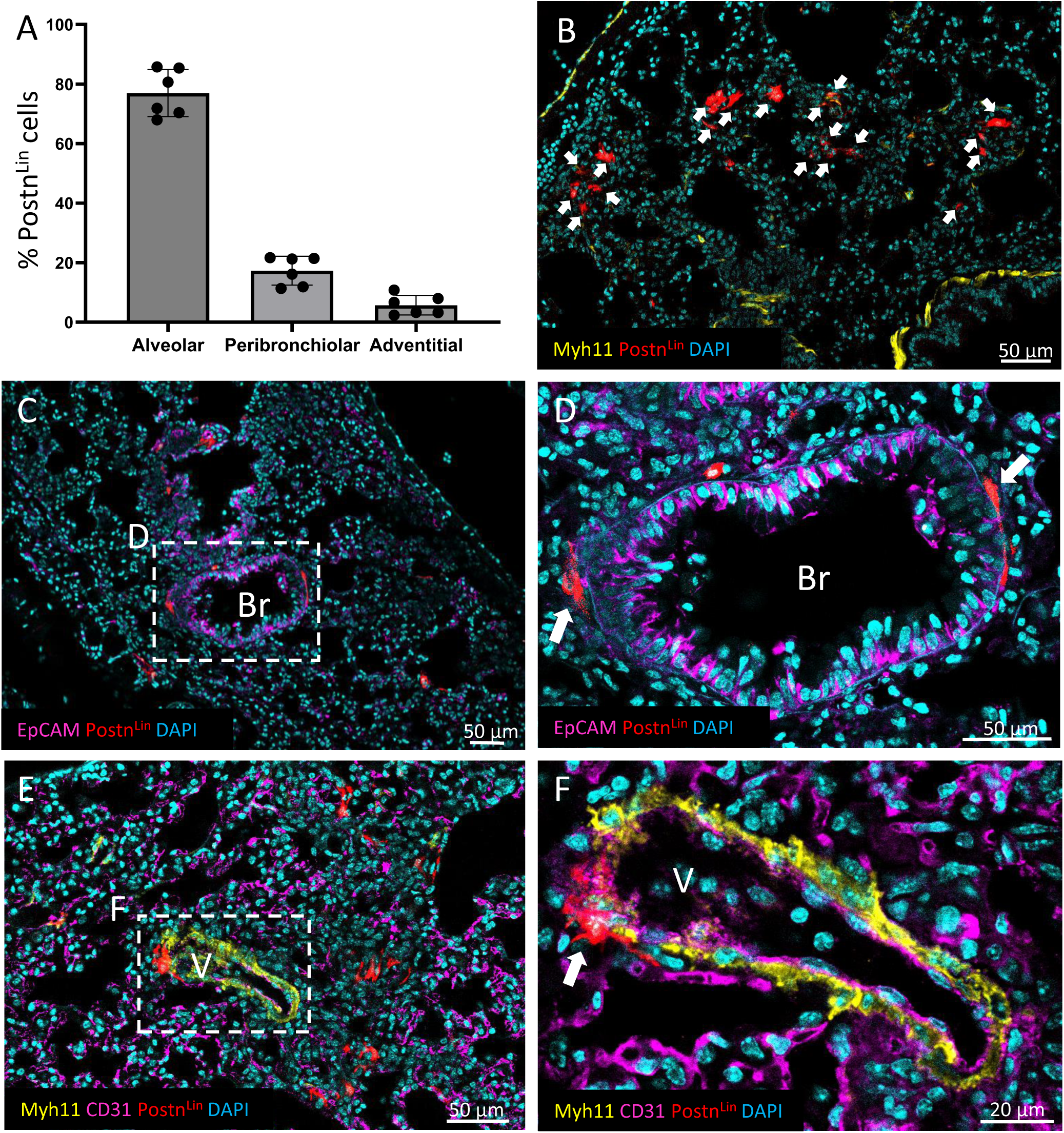
Postn^Lin^ cells are predominantly in the alveolar interstitium. (A) Quantification of the proportion of Postn^Lin^ cells in alveolar, peribronchiolar, and adventitial locations by immunohistochemistry. (B) Representative image of Postn^Lin^ cells in alveolar spaces, highlighted with white arrows. (C) Low-power and (D) high-power images of peribronchiolar (Br) localization: arrows point to Postn^Lin^ cells located beneath the bronchiolar epithelium stained with an antibody against the epithelial marker EpCAM. (E) Low-power and (F) high-power images of Postn^Lin^ cells in an adventitial location surrounding a vessel (V). Arrow indicates Postn^Lin^ cells underlying the perivascular smooth muscle. Endothelial cells are highlighted using an antibody against CD31, whereas smooth muscle cells are stained with an antibody against Myh11.

### Postn^Lin^ cells induced by *A. fumigatus* belong to the fibroblast population

Platelet-derived growth factor α (Pdgfrα) is expressed by most tissue resident fibroblasts and is widely used as a marker for that cell type, including for those found in the lung^11,20^. We therefore used a Pdgfrα lineage (Pdgfrα^Lin^) tracing mouse to compare the morphology of Pdgfrα^Lin^ resident pulmonary fibroblasts to that of *A. fumigatus*-induced Postn^Lin^ fibroblasts. Resident Pdgfrα^Lin^ fibroblasts displayed a thin, elongated morphology and were widely distributed throughout the lung in the same alveolar, peribronchial and adventitial locations as Postn^Lin^ cells (Fig. S4A-D). Postn^Lin^ fibroblasts that localized to peribronchial and adventitial locations showed a similar morphology, but those that were found clustered among regions of inflammation in the alveoli displayed a more hypertrophic and dendritic morphology with numerous branched extensions (Fig. S4E-H). To determine whether Postn^Lin^ cells express the Pdgfrα fibroblast marker, the Postn^Lin^ mice were crossed with a Pdgfrα-GFP mouse in which the endogenous Pdgfrα promoter constitutively drives expression of a histone H2B-eGFP fusion protein^21,22^. The resulting Postn^Lin^/Pdgfrα-GFP strain was inoculated with 3 x 10^7^ conidia and examination of lung sections on day 7 post-inoculation revealed a subset of Postn^Lin^ fibroblasts that also expressed Pdgfrα (Fig. 4A and B, orange arrows). Quantification by flow cytometry showed that approximately 60% of the tdTomato+ cells coexpressed Pdgfrα (Fig. 4C), similar to what has been previously reported for Postn^Lin^ cells in the heart ^11,16,22^.

**Figure 4.**
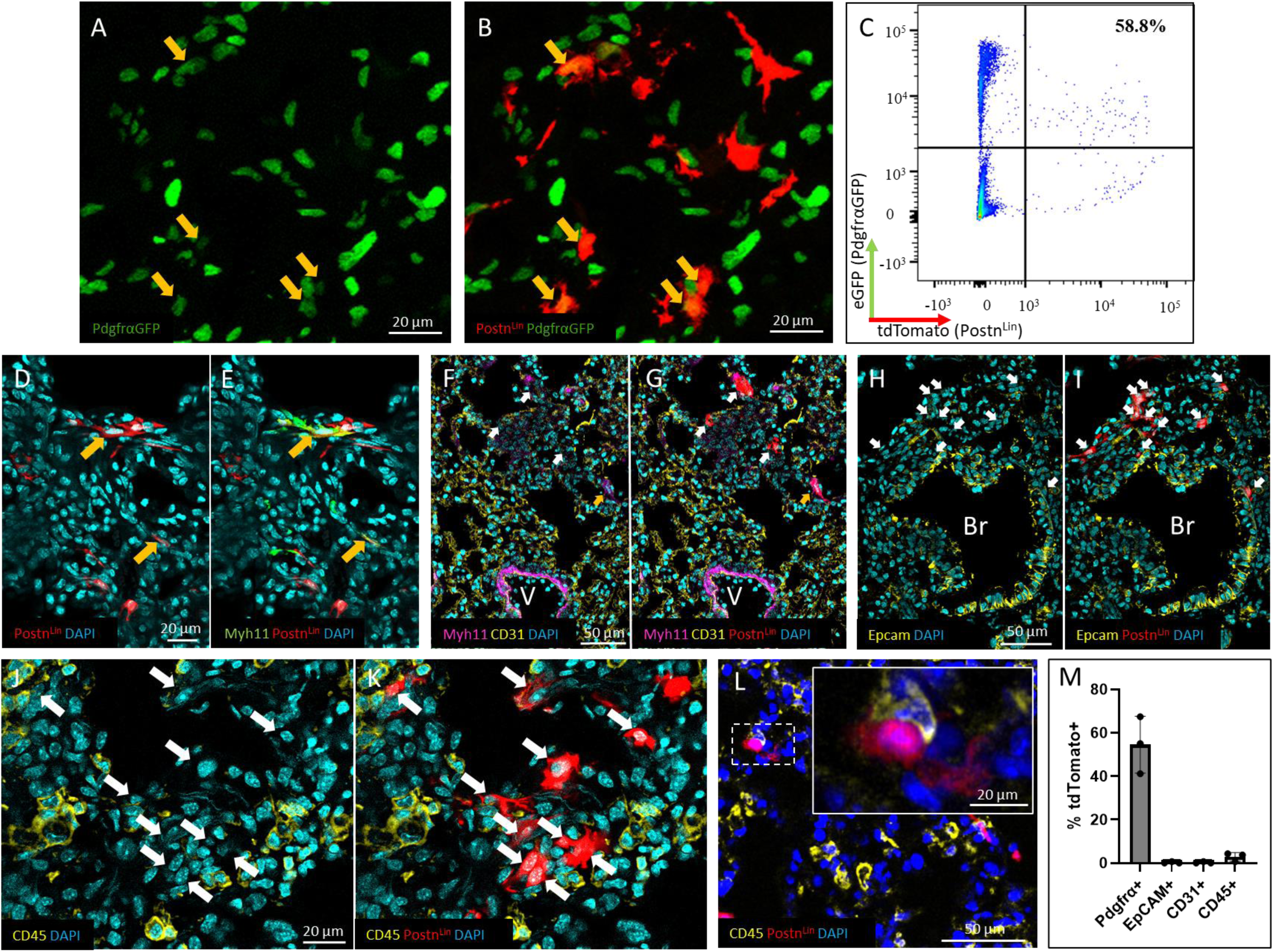
Postn^Lin^ cells belong to the fibroblast population. (A & B) Images display colabeling of Pdgfrα-GFP (green nuclei) with *A. fumigatus*-induced Postn^Lin^ cells (red cells), highlighted by yellow arrows. (C) Representative flow cytometry plot showing tdTomato+ cells (rightward scatter) against Pdgfrα cells (upwards scatter). The percentage of Pdgfrα+ cells within the tdTomato+ population is depicted in the upper right quadrant. (D & E) Images show Postn^Lin^ cells (red) and Myh11-labeled cells, with DAPI- stained nuclei in blue. Yellow arrows show costaining. (F-K) Representative images showing the lack of colocalization between Postn^Lin^ cells (red) and antibody staining against the endothelial marker CD31 (F&G), the epithelial marker EpCAM (H&I), and the hematopoietic marker CD45 (J-L). The color coding is shown at the base of the figure. White arrows highlight the location of Postn^Lin^ cells to facilitate the comparison of colocalization. The inset in panel L represents an example of how Postn^Lin^ cells are often observed in close contact with hematopoietic cells. (M) Flow cytometric quantification of tdTomato+ cells colocabeled with markers for fibroblasts (Pdgfrα), smooth muscle (Myh11), epithelial cells (EpCAM), endothelial cells (CD31), and hematopoietic cells (CD45). Values represent the mean ± SD from 3 biological replicates.

We also found that approximately 25% of the tdTomato+ cells costained with smooth muscle myosin heavy chain 11 (Myh11, Fig. 4D and E), a marker that is expressed by smooth muscle cells as well as a subset of alveolar fibroblasts that have differentiated to a contractile myofibroblast phenotype^11,23^ (Fig. 4D and E, colocalization shown by orange arrows). Most of the observed Myh11+/tdTomato+ colocalization was in alveolar regions where smooth muscle is absent (Fig. 4D and E), suggesting that the majority of the double-labeled cells represent contractile myofibroblasts. By contrast, tdTomato+ cells did not costain with markers for endothelial cells (CD31, Fig. 4F, G) or epithelial cells (EpCAM, Fig. 4H, I) by immunohistochemistry. Similarly, tdTomato+ cells did not colabel with the hematopoietic marker CD45 by immunohistochemistry (Fig. 4J– L). However, CD45+ cells were often observed in close proximity to tdTomato+ cells (Fig. 4L, inset). The absence of colabeling with non-stromal populations was confirmed by flow cytometry, which revealed minimal numbers of tdTomato+/CD45+ cells (Fig. 4M and Fig. S5).

We conclude that Postn^Lin^ cells induced by the presence of *A. fumigatus* in the lung belong to the pulmonary fibroblast population and that they show the morphology of an activated fibroblast state that is distinct from that of resident Pdgfrα^Lin^ fibroblasts.

### Targeted ablation of the Postn^Lin^ lineage exacerbates the severity of invasive aspergillosis

To determine the impact of Postn^Lin^ cells on the pathogenesis of IPA, we used a genetically modified mouse line for conditional cell ablation that employs inducible expression of diphtheria toxin subunit A (DTA) gene under the control of a Cre recombinase^24^. Specifically, a TAM-inducible Cre recombinase (Postn-MerCreMer) was targeted into one Postn allele and crossed with a strain that expresses DTA in response to Cre activity (Fig. 5A). In this mouse line, fibroblasts that activate the Postn promoter will express DTA, thereby killing the cells and preventing fibroblast activity and downstream responses. To induce IPA, we modified a triamcinolone-induced immunosuppression protocol^25^. Mice were inoculated with 1 x 10^6^ conidia and the host-pathogen interaction was allowed to occur for 5 hours in the absence of immunosuppression. The mice were then immunosuppressed with a single dose of triamcinolone acetonide, and the progression of IPA was monitored over time (Fig 5B). The mortality rate of infected DTA mice was accelerated relative to infected control mice (Fig. 5C) and was associated with visible evidence of hemoptysis. Dissected lungs from the infected DTA mice showed gross evidence of pulmonary hemorrhage (Fig. 5D), and histopathological analysis revealed focal areas of alveolar hemorrhage with extensive fungal angioinvasion (Fig. 5E). Hemoptysis and alveolar hemorrhage were notably absent from infected control mice. In the absence of immunosuppression, there was no death of DTA mice following an OP challenge with *A. fumigatus* conidia, even at a higher inoculum of 3 x 10^7^ conidia (data not shown). This suggests that the loss of Postn^Lin^ cells in the context of an invasive infection increases susceptibility to *A. fumigatus*-induced vascular damage.

**Figure 5.**
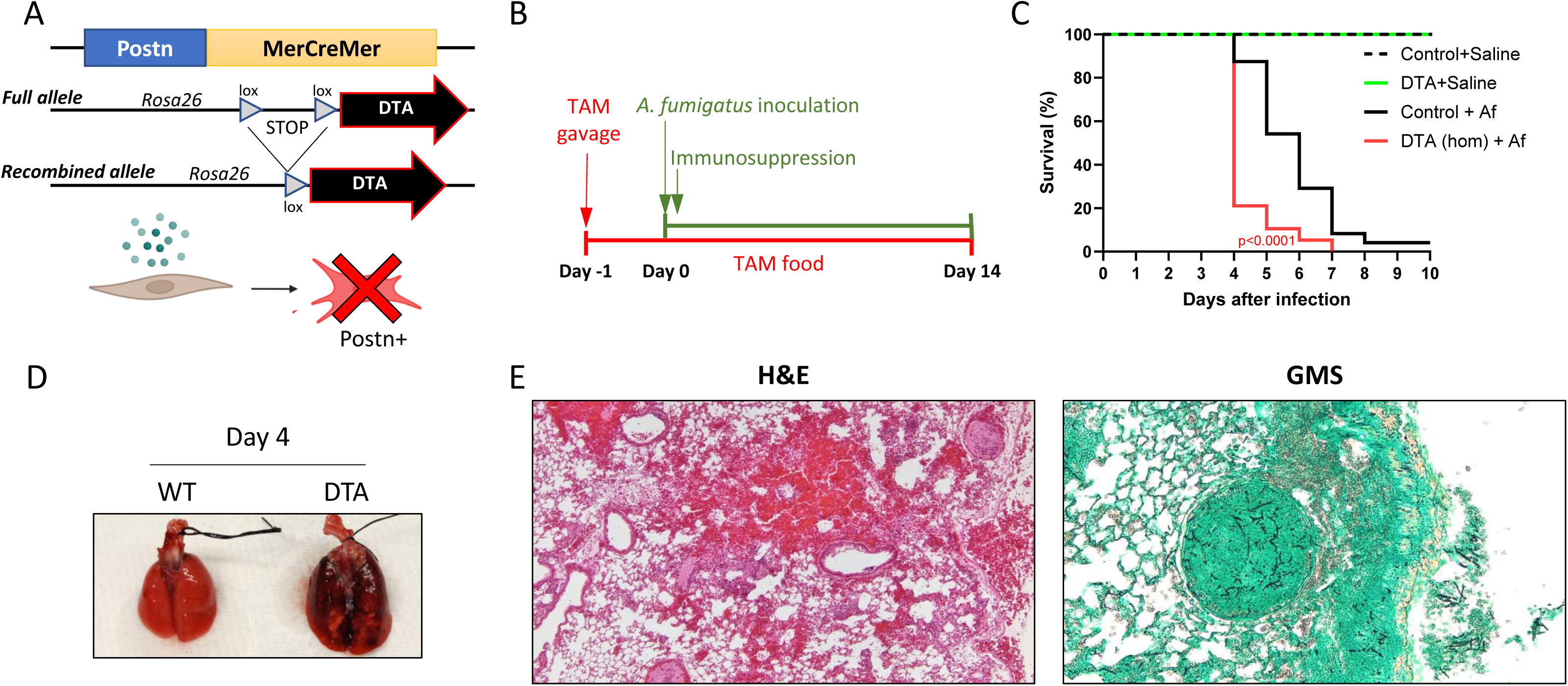
Targeted ablation of Postn^Lin^ cells accelerates the progression of IPA. (A) Schematic of the TAM-inducible DTA ablation model. The Cre recombinase, inserted into the Postn locus, is controlled by a TAM-inducible estrogen receptor (MerCreMer). Upon Postn promoter activation, TAM-induced recombination of the DTA gene results in DTA toxin expression from the rosa26 locus, preventing fibroblast activation and expansion. (B) Schematic representation of the experimental approach: Mice were inoculated OP with 1 x 10^6^ conidia, followed by immunosuppression with triamcinolone acetonide 5 hours post-infection. TAM was administered via gavage one day before fungal inoculation and maintained through a TAM-infused chewing diet for the duration of the experiment. (C) Percent survival of immunosuppressed mice infected with *A. fumigatus* (n=19; P, < 0.0001, log rank test). (D) Gross appearance of lungs from infected DTA and control mice on day 4 post-infection. (E) Representative histopathologic images showing focal alveolar hemorrhage and angioinvasion in the *A. fumigatus*-infected DTA mice by H&E and GMS staining, respectively.

### Postn^Lin^ cells express a unique transcriptional profile of matrix modification compared to resident pulmonary fibroblasts

To assess how the *A. fumigatus*-host interaction impacts gene expression in the stromal population, we performed bulk RNA-seq on *A. fumigatus*-induced Postn^Lin^ fibroblasts, using resident Pdgfrα-GFP fibroblasts as the comparator. Immunocompetent Postn^Lin^ mice were administered a single inoculum of *A. fumigatus* conidia to induce fibroblast activation and Pdgfrα-GFP mice were given a mock inoculation with saline. The induced Postn^Lin^ cells and the resident Pdgfrα-GFP fibroblasts were then isolated on day 7 post-inoculation by sorting tdTomato+ or GFP+ cells that were negative for vascular (CD31), epithelial (Epcam), and hematopoietic (CD45) markers (see Fig. S6 for sorting strategy). Total RNA was extracted from the two populations and bulk RNA-Seq was performed to compare the transcriptional profile of *A. fumigatus*-induced Postn^Lin^ cells to that of Pdgfrα+ resident fibroblasts.

A total of 1037 differentially expressed genes (DEGs) were identified using a statistical cutoff of a Bonferroni-adjusted P-value ≤ 0.05 and a fold change greater than ± 1.5 (Fig. 6A; Table S1). Among these DEGs, 552 were upregulated in the Postn^Lin^ population while 485 were downregulated, which are highlighted in the volcano plot shown in Fig. 6A. Genes involved in ECM organization, cell adhesion, and angiogenesis were among the top 10 enriched Gene Ontology (GO) terms in both the upregulated and downregulated categories (Fig. 6B and C), demonstrating that the tissue remodeling activity of Postn^Lin^ fibroblasts is distinct from that of resident pulmonary fibroblasts. Downregulated genes were enriched in GO categories associated with signaling pathways, metabolism, gene expression, cell differentiation and proliferation (Fig. 5C; Table S3) reflecting the comprehensive transcriptional rewiring that is necessary for resting fibroblasts to transition to an activated phenotype. Several of the genes that were downregulated in response to *A. fumigatus* represent markers of resident pulmonary fibroblasts, including Pdgfrα, Tcf21, and Scube2^26^ (Fig. 5D), suggesting that pathogen exposure induces a shift away from the gene signature of resident fibroblasts.

**Figure 6.**
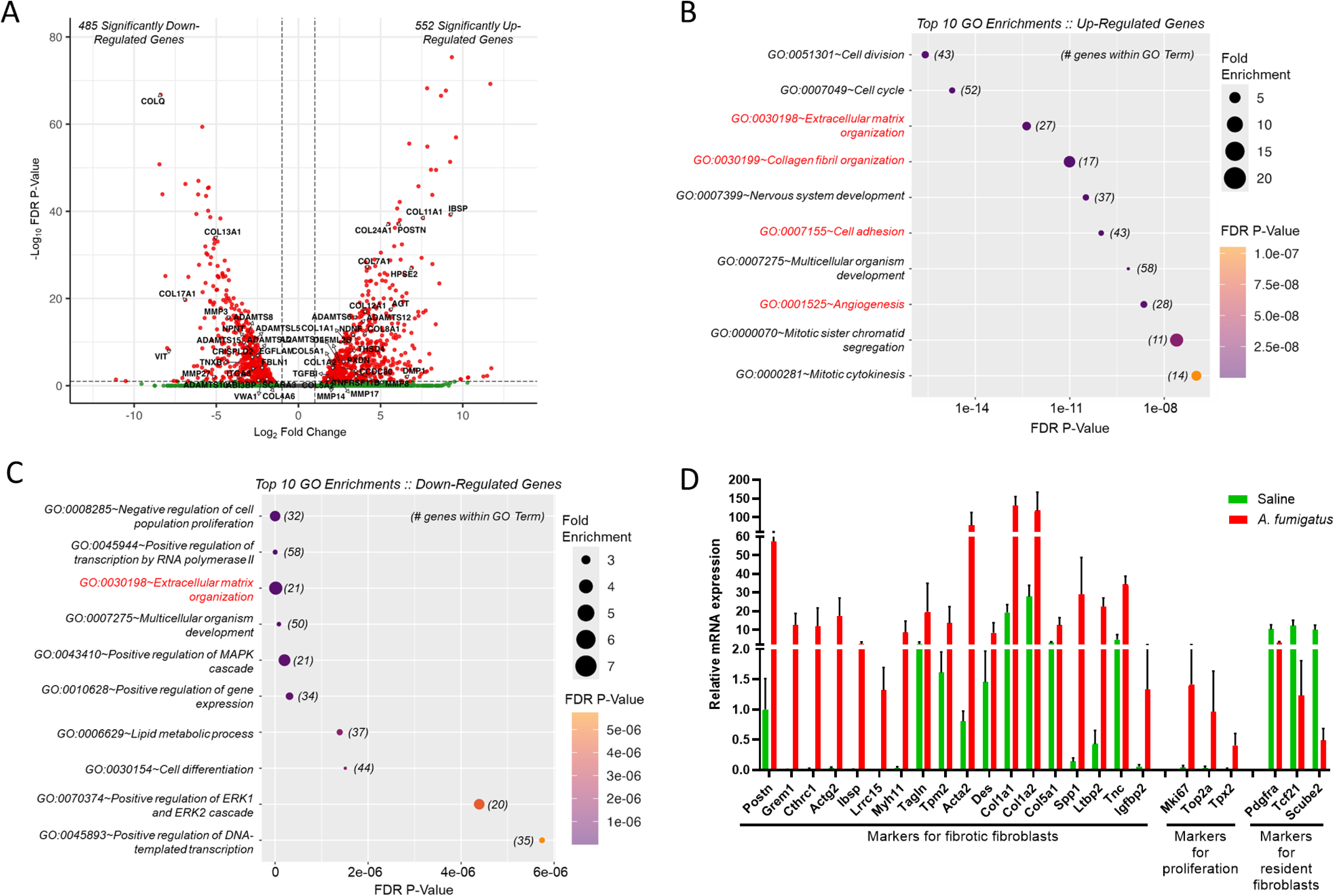
PostnLin cells express a unique transcriptional profile of ECM modification compared to resident pulmonary fibroblasts. Induced Postn^Lin^ cells and resident Pdgfrα^Lin^ fibroblasts were isolated by flow cytometry as described in methods and total RNA was sequenced. (A) Volcano plot shows 485 down-regulated and 552 up- regulated genes after *A. fumigatus* infection. Genes related to extracellular matrix organization (GO:0030198) are highlighted. (B & C) Top 10 *A. fumigatus*-dependent GO Terms for significantly (B) upregulated and (C) downregulated genes, ranked by P-value and fold enrichment. Matrix remodeling terms are highlighted in red. (D) Expression comparison of selected differentially expressed genes between *A. fumigatus*-induced Postn^Lin^ cells and mock-treated Pdgfrα^Lin^ cells. Values represent the means ± SD from 4 biological replicates.

Among the top upregulated DEGs were those involved in fibroblast activation and fibrosis (selected genes are highlighted in Fig. 5D). These include: Grem1 (Gremlin1), a bone morphogenic protein agonist involved in ECM regulation and fibrosis^27,28^; Cthrc1 (collagen triple helix repeat containing 1), a secreted ECM component implicated as the major effector of fibrotic processes in the injured lung^26^; Actg2 (actin gamma 2), linked to fibroblast activation in the lung^29^; IBSP (integrin binding sialoprotein (IBSP), identified in fibrotic lungs^30^; and Lrrc15 (leucine-rich repeat containing 15), associated with myofibroblast populations^31–33^. Additional DEGs previously reported to be associated with fibrotic fibroblast activity included Myh11, Tagln, Tpm2, Acta2, Des, Col1a1, Col1a2, Col5a1, Spp1, Ltbp2, Tnc and Igfbp2^26,29^ (Fig. 5D). These results suggest that rewiring of fibroblast gene expression to modify the ECM is central to confronting an infectious threat from *A. fumigatus*, consistent with our observation of collagen accumulation within inflammatory foci in the lung induced by *A. fumigatus* (Fig. 1C).

### Pulmonary fibroblast sub-populations with enhanced matrix stabilizing and immunomodulatory characteristics emerge following a challenge with *A. fumigatus*

Our bulk RNA-seq analysis was performed at day 7 post-inoculation due to peak accumulation of tdTomato+ fibroblasts at that point (Fig. 2D and E), revealing a predominant gene expression signature of matrix-modifying activity. To determine whether additional fibroblast heterogeneity emerges at earlier stages of the host-fungal interaction and to delineate other potentially important subsets of fibroblasts that could be masked by the major populations, we used scRNA-sequencing (scRNA-seq). Immunocompetent mice were exposed to either a single inoculum of conidia or a mock inoculation with saline. The stromal cell population was then isolated from total lung cells on days 1 and 7 post-inoculation by flow cytometry, excluding the non-stromal population based on expression of CD31, CD45 and EpCAM, consistent with the bulk RNA-seq analysis (Fig 7, see Fig. S6 for sorting strategy). The resulting stromal cell population was then subjected to a single-cell transcriptome analysis.

**Figure 7.**
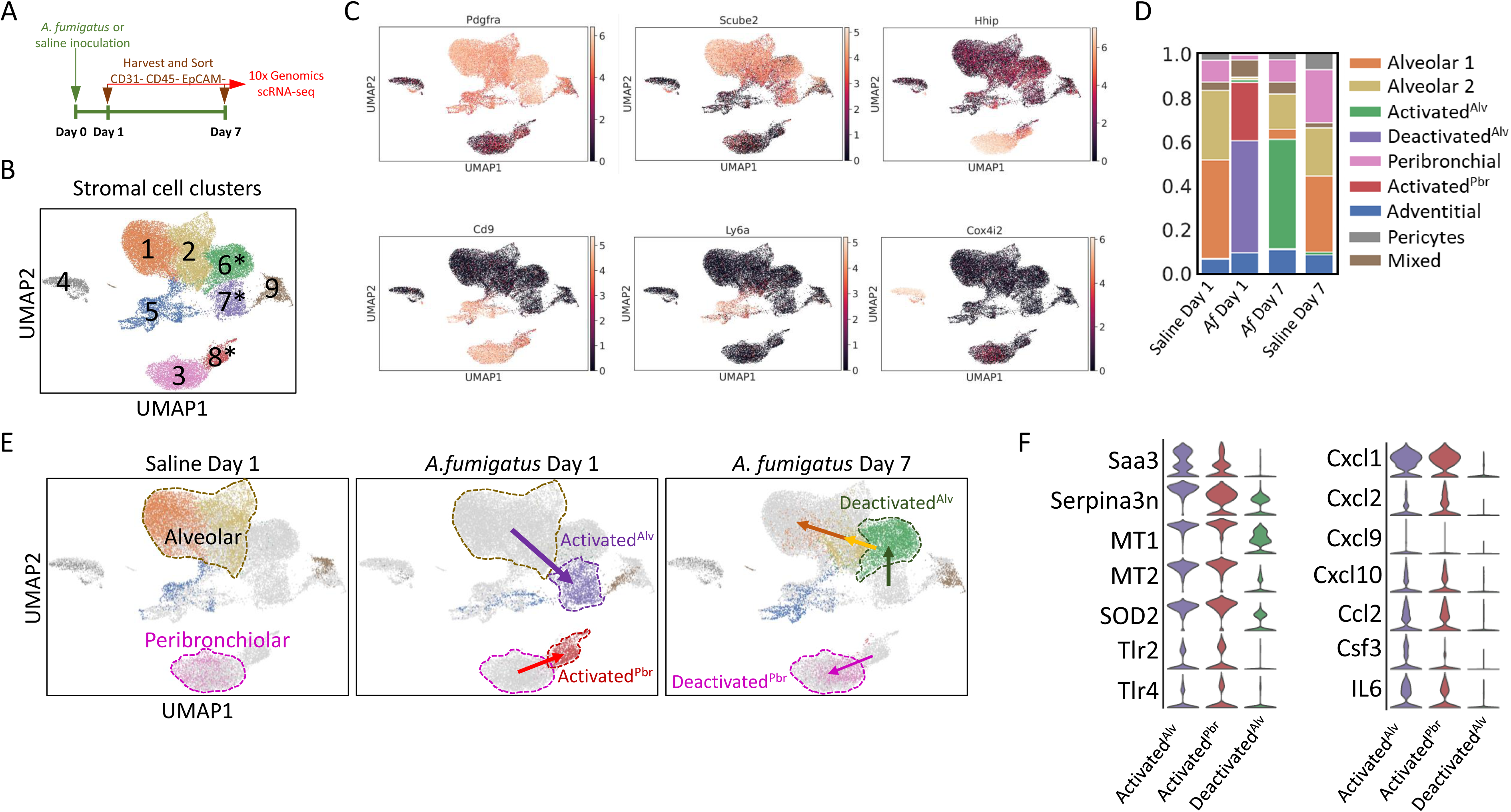
Pulmonary fibroblast sub-populations with enhanced matrix stabilizing and immunomodulatory characteristics emerge following a challenge with *A. fumigatus*. (A) Schematic representation of the experimental approach: Mice were inoculated OP with 3 x 107 conidia, and lungs were harvested and digested at day 1 and 7 post-inoculation. Stromal cells were sorted by excluding CD31+, CD45+ and EpCAM+ populations. (B) Uniform Manifold Approximation and Projection (UMAP) plot showing 9 stromal cell clusters on days 1 and 7 after pulmonary challenge with A. fumigatus conidia or mock inoculation with saline. Asterisks highlight clusters that are unique for the *A. fumigatus*-inoculated mice. (C) UMAP plots illustrate the expression of the indicated genes across the different clusters. The expression of these genes correlates with fibroblasts located at different anatomical sites as has been previously shown19: Pdgfrα+ and Scube2+ for alveolar fibroblasts, Hhip+ and CD9+ for peribronchial fibroblasts, Ly6a+ and CD9+ for adventitial fibroblasts, and Cox4i2 for pericytes. (D) Fraction bar plot representing the relative proportion of each cluster in the four experimental groups. (E) UMAP plots highlight clusters that uniquely emerge on day 1 and 7 post-inoculation with saline or *A. fumigatus*. Arrows indicate the proposed trajectory based on gene expression analysis. (F) Violin plots representing inflammatory genes induced exclusively in the *A. fumigatus* induced clusters. Day 7 post-inoculation cluster reflects partial recovery to a non-inflammatory state.

Cell typing was performed using unsupervised clustering and a reference scRNA- seq dataset of pulmonary fibroblast populations^19^. This approach identified 9 distinct fibroblast populations (Fig. 7B), with clusters 1 and 2 classified as alveolar fibroblasts, cluster 3 as peribronchial fibroblasts, cluster 4 as pericytes, cluster 5 as adventitial fibroblasts, clusters 6-8 as *A. fumigatus*-induced clusters (highlighted by the asterisks), and cluster 9 as an indistinct population of fibroblasts with mixed characteristics. Representative genes used for cell typing are shown in Fig. 7C. As expected, Pdgfrα+ cells were distributed across multiple clusters, consistent with the experimental enrichment of a stromal fibroblast population. Scube2+ cells, representing an alveolar fibroblast phenotype, were distributed across 4 clusters (Fig. 7B: clusters 1, 2, 6 and 7); Hhip+/Cd9+ cells, indicative of peribronchial fibroblasts, were present in 2 clusters (Fig. 7B: clusters 3 and 8); Ly6a+/Cd9+ cells, indicating adventitial fibroblasts were located in cluster 5 (Fig.7B); and cluster 4 was marked by the pericyte marker Cox4i2 (Fig.7B). Cells in the alveolar and peribronchial fibroblast clusters in the saline-inoculated mice (Fig. 7D and E, left panel) reduced in number in response to *A. fumigatus* on day 1 post-infection (Fig. 7D and E, middle panel), coinciding with the appearance of an activated alveolar fibroblast cluster (Activated^Alv^) and an activated peribronchial fibroblast cluster (Activated^Pbr^, purple and red clusters, respectively).

Analysis of the differentially expressed genes (DEGs) contained within the various activated fibroblasts identified unique and overlapping genes that fell into two notable categories: ECM synthesis and immune modulation (Table S4). In the ECM category, both activated clusters were enriched for genes involved in the formation of matrix structural fibers, encoding 9 different types of collagen, as well as elastin, fibrillin, and three secreted lysyl oxidase enzymes with roles in the formation of collagen and elastin crosslinks that stabilize the ECM^34–36^ (Table S4). In addition, several genes encoding integrins that represent the main cellular receptors used to interact with proteins in the ECM were shared between both clusters. In many organs, the TGFβ regulatory system plays a central role in driving *de novo* collagen synthesis^34^. This fibrogenic function of TGFβ is transduced by the cytosolic signaling protein Smad3 and is tightly regulated by latent TGF-β binding protein 1 (ltbp1), which sequesters TGFβ in the ECM. The observed enrichment of TGFβ, ltbp1, and Smad3 in the activated alveolar fibroblast cluster is therefore consistent with a predominant fibrogenic role for these cells (Table S4). This cluster was also remarkable for the presence of 16 DEGs encoding matrix-modifying proteins, indicating that its component cells were actively engaged in remodeling the ECM^37^. The expression of α-smooth muscle actin (Acta2) was notably upregulated in the activated peribronchial cluster, suggesting that some of these cells had differentiated to a contractile myofibroblast phenotype^38^.

The second group of DEGs in the activated clusters were associated with immune modulatory activity (Table S4), particularly in the activated alveolar fibroblast population. These included the cytokines IL-6, IL-10, and IL-11, as well as multiple cytokines that are members of the TNF, TGF-β, and BMP superfamilies (Table S4). The presence of TGF-β is particularly significant because of its dual role in regulating immune responses as well as fibrogenesis^34^. Although immune defense against *A. fumigatus* relies heavily on tissue-resident alveolar macrophages and dendritic cells, additional inflammatory cells such as neutrophils, inflammatory monocytes, and monocyte-derived dendritic cells are recruited to the site of infection as needed^39^. The signal that brings these cells into the lung involves the secretion of chemokines^39^. Interestingly, both activated clusters were enriched for 11 of these chemokines, many of which have established roles in host defense against *A. fumigatus* and other IFIs^7^^40–46^. Upon arrival in the lung, the functional activity of recruited myeloid cells is regulated by cytokines such CSF1 (macrophage colony-stimulating factor) and CSF3 (granulocyte colony stimulating factor), both of which were enriched in the activated fibroblasts clusters. Immune cell activation is also accomplished by lipid mediators such as the prostaglandins^47^. DEGs involved in prostaglandin synthesis were present in both activated clusters, indicating that activated fibroblasts share cytokine- and eicosanoid-mediated signaling to modulate the immune system. In addition to cytokines, these activated alveolar fibroblasts were enriched for receptors for 25 different cytokine and prostaglandin ligands, indicating that the communication between fibroblasts and the immune system is bidirectional.

Pattern recognition receptors (PRRs) are a class of receptors on cells of the innate immune system that are on the front lines of innate immune cell recognition, binding to pathogen-associated molecular patterns (PAMPs) on microbial surfaces and triggering downstream immune activity^39^. Both activated fibroblasts clusters were enriched for PRRs, suggesting that fibroblast activation would enhance the ability of these cells to recognize microbial pathogens. PRRs identified in the clusters included the cell surface Toll-like receptors TLR-2 and TLR-4, the intracellular sensors NOD1 and NLRC3, and the secreted PRRs that include the galectin family of lectins and pentraxin-3 (PTX3)^48–52^.

By day 7 post-inoculation, the number of cells in the activated alveolar fibroblast cluster was dramatically reduced, coinciding with the appearance of another *A. fumigatus*- specific cluster that included DEGs encoding resident fibroblast markers Tcf21 and Pdgfrα, as well as the alveolar fibroblast markers Scube2, Npnt, Ces1d and Inmt (Fig. 7D and E, right panel, green cluster and Table 1). This suggests that activated alveolar fibroblasts are beginning to deactivate, indicating a progressive return to a resident alveolar fibroblast state (Fig. 7E, arrows highlight the proposed trajectory). Similarly, by day 7 the activated peribronchial fibroblast population was no longer evident (Fig. 7D and E), while the resident peribronchial fibroblast cluster was reconstituting (Fig. 7E, arrows highlight the proposed trajectory). A representative set of inflammatory genes previously upregulated in the activated alveolar and activated peribronchiolar clusters was markedly downregulated in the deactivated alveolar fibroblast cluster (Fig. 7F), indicating progressive transition to a quiescent state. The return of activated fibroblast clusters to a resident phenotype is consistent with the observed histologic evidence of resolving inflammatory foci by day 7 post-inoculation with *A. fumigatus* (Fig. 1).

**Table 1.**
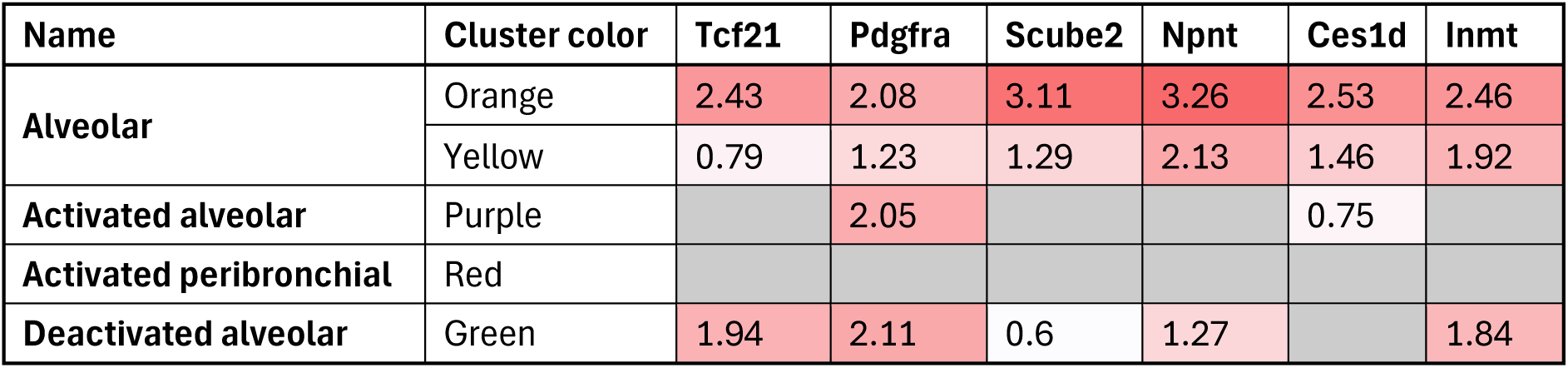
Resident alveolar markers expressed by cells in the indicated clusters. Values represent the log fold change.

We conclude that exposure to an infectious threat with *A. fumigatus* triggers pulmonary stromal cells to differentiate into an emergent population of ECM-producing and immunomodulatory fibroblasts, which progressively return to a resident state as the infection resolves.

## DISCUSSION

In nature, *A. fumigatus* secretes hydrolytic enzymes for the purpose of breaking down organic material into easily metabolizable substrates. Consequently, one of the shared features of all human pulmonary infections caused by this fungus is tissue injury. Here, we provide evidence that the stromal population of the lung is an integral component of the response to fungus-mediated damage. Specifically, we demonstrate that newly synthesized collagen fibers are deposited within areas of pulmonary inflammation induced by the inhalation of *A. fumigatus* conidia (Fig.1). By employing a lineage-tracing approach that uses Postn gene expression as the marker for fibroblast activation, we show that Postn^Lin^ fibroblasts emerge in the vicinity of *A. fumigatus*-induced inflammatory foci, acquiring an enlarged dendritic morphology that would increase fibroblast surface area. (Fig. S4). This morphology contrasts the thin elongated appearance of resident Pdgfrα^Lin^ fibroblast population in uninjured lung tissue. Immunohistochemical and flow cytometric analyses confirmed that *A. fumigatus*-induced Postn^Lin^ cells are part of the stromal population, and that most of them localize to the alveolar interstitium, with a few residing beneath the epithelium of the conducting airways. Immunohistochemical and flow cytometric analyses confirmed that *A. fumigatus*-induced Postn^Lin^ cells are part of the stromal population, and that most of them localize to the alveolar interstitium, with a few residing beneath the epithelium of the conducting airways.

Notably, a comparable level of activation was observed following inoculation with heat-killed conidia, suggesting that the surface of a dormant inhaled conidium, coated with a hydrophobic layer of rodlet proteins^39^, is sufficient to induce this response.

Using a conditional cell ablation model, we demonstrate that the absence of Postn^Lin^ cells during IPA is associated with pulmonary hemorrhage and fungal angioinvasion combined with an accelerated mortality rate (Fig. 5). Most of these Postn^Lin^ cells are Pdgfrα+, suggesting that they arise from the resident fibroblast populations in the lung. However, pericytes, which are fibroblast-like cells located on the abluminal surface of the microvasculature^53^, have also been reported to express Postn^26^. This suggests that the cell ablation model used here could potentially target both resident pericytes and activated fibroblast subsets. Since pericytes support microvascular homeostasis^53^, and *A. fumigatus* proteases degrade collagen types I and III, elastin, and fibronectin in the ECM^54,55^, we speculate that ablating the Postn^Lin^ lineage diminishes microvascular stability and reduces the ability of fibroblasts to repair proteolytic damage to the ECM, resulting in an overall weakening of pulmonary tissue that predisposes to fungal invasion and alveolar hemorrhage in the course of infection.

We employed scRNA-seq to gain insight into the emergence of fibroblast heterogeneity in response to *A. fumigatus*. One of the most striking findings was the transient appearance of two clusters of cells that were unique to mice inoculated with *A. fumigatus*. These clusters were enriched in collagen fiber synthetic genes, encoding both collagen structural proteins and the oxidase and hydroxylase enzymes involved in their synthesis (P4ha2, P4ha3, Plod2, Lox, Loxl1, Loxl2). Col1a1 is the most abundant collagen and was enriched in resident alveolar fibroblasts. However, it was not enriched in the activated fibroblast clusters, which instead expressed a unique array of collagen genes (Table S4). This suggests that *A. fumigatus* triggers stromal fibroblasts to switch to the synthesis of a different set of collagens that are necessary for expansion and/or repair of the ECM. The activated alveolar fibroblast population was notably enriched in elastin and fibrillin genes, which synergize to confer both tensile strength and elasticity to lung tissue^56^. In addition, fibrillin is indirectly involved in the regulation of TGFβ activation and fibrogenesis through its interaction with latent TGFβ binding proteins^57^. A previous scRNA-seq analysis of collagen-producing cells in the lung identified a subpopulation that emerges in the context of fibrotic lung disease and is characterized by expression of the collagen triple helix repeat containing-1 gene (Cthrc1)^26^. Interestingly, Cthrc1 was identified as a top DEG in our bulk RNA sequencing of Postn^Lin^ cells, but it was not detected in the activated clusters identified through scRNA-seq. This discrepancy may be due to the bulk RNA-seq analysis enriching for Postn^Lin^ cells that have high levels of Cthrc1 (Fig. 6D), while Cthrc1+ cells are less apparent in the larger stromal cell populations examined by scRNA-seq analysis. We speculate that fibroblasts in healthy lungs are equipped to respond to the daily inhalation of conidia without triggering major expansion and activation of a population of Cthrc1+ fibroblasts that could lead to pathological fibrosis.

Emerging data shows that fibroblasts have the capacity to influence inflammatory processes^58^. Our data support this, showing that fibroblasts activated by *A. fumigatus* upregulate the expression of cytokines and chemokines, most of which are pro-inflammatory. In addition, the enrichment of numerous cytokine receptors (Table S4) indicates that fibroblasts and immune cells engage in reciprocal communication to influence the outcome of infection. Our studies reveal that pulmonary fibroblasts may also play a direct role in pathogen recognition since the expression of cell surface, intracellular, and secreted PRRs is enhanced during activation. PTX3 is a secreted PRR that is particularly relevant to host defense against *A. fumigatus* since it functions as an opsonin with antifungal activity, and mice lacking PTX3 have increased susceptibility to IPA that can be countered by exogenous administration of PTX3^51,52,59^. In addition, we observed upregulation of the cell surface PRRs TLR-2 and TLR-3, which recognize swollen conidia, germlings, and hyphal forms of *A. fumigatus*^39^. Although these immune functions are relevant to immune protection against *A. fumigatus*, the mechanisms involved have broad specificity that would also be protective against different pathogens. It will be of considerable interest in future studies to determine whether enhanced immunomodulatory activity is a generalized response of fibroblasts to infection or whether fibroblasts can tailor their response to a specific pathogen.

Despite these major changes in gene expression, the activated fibroblast clusters were not permanent; both showed evidence of deactivation coinciding with fungal clearance and the progressive resolution of inflammatory cell foci in the lung (Fig. 1 and S1). Taken together, these studies reveal that fibroblasts are active participants in lung protection through the transitory induction of immunomodulatory and ECM-preserving functions that mitigate the tissue damage associated with a pulmonary infection with *A. fumigatus*. These findings may offer a foundation for future targeted therapeutic strategies aimed at bolstering host defenses against *A. fumigatus*-induced pulmonary pathology.

## METHODS

### Fungal strains and growth conditions

The *A. fumigatus* clinical isolate CEA10^60,61^ was used throughout the study, while the clinical isolates Af293 and H237^62^ were used where indicated. Conidia were harvested before each experiment from mycelia grown for 3 days at 37⁰C on solid *Aspergillus* minimal medium (AMM) (1% [wt/vol] D-glucose, 1% [vol/vol] NH4 tartrate, 2% [vol/vol] salt solution [2.6% {wt/vol} KCl, 2.6% {wt/vol} MgSO4 heptahydrate, 7.6% {wt/vol} KH2PO4, and 5% {vol/vol} trace element solution]), then washed and resuspended in saline. When required, conidia were inactivated by heat shock at 65⁰C for 30 min. The effectiveness of heat inactivation was confirmed by plating onto the center of a plate of YG medium (0.5 yeast extract, 2% glucose) plates, and confirming no surviving CFUs after 4 days of incubation at 37⁰C. Mice were inoculated as described in *Animal Procedures*. Colony formation units (CFUs) were quantified by spreading 100 μL of serially diluted lung homogenates onto Becton Dickinson (BD) inhibitory mold agar (IMA). The plates were incubated for 16-24 h at 37⁰C. At least 3 biological replicates were analyzed per condition.

### Mouse lines

Rosa26-tdTomato (B6.Cg-*Gt(rosa)26Sor^tm14(CAG-tdTomato)Hze^*/J; JAX Strain #:007914), Rosa26-DTA (B6.129S6(Cg)-*Gt(ROSA))26Sor^tm1(DTAJpmb^*/J; JAX Strain #:032087), Pdgfrα-CreERT2 (B6.129S-*Pdgfra^tm1.1(cre/ERT2)Blh^*/J; JAX Strain #:032770), and Pdgfrα-eGFP (B6.129S4-*Pdgfra^tm11(EGFP)Sor^*/J; JAX Strain #:007669) mice were purchased from The Jackson Laboratory. PostnMerCreMer (Postn^MCM^) (B6.129S-Postntm2.1(cre/Esr1*)Jmol/J; JAX Strain #:029645) mice were generated as previously described^16^. Postn^MCM^ and Pdgfrα-Cre mice were crossed with a Rosa26-tdTomato mouse line to enable reporter expression upon TAM-induced Cre recombination in Postn- or Pdgfra-expressing cells. Similarly, Postn^MCM^ mice were crossed with Rosa26-DTA mice to induce specific ablation of Postn-expressing cells in the presence of TAM. Heterozygous Postn^MCM^ and Pdgfrα-Cre mice were utilized for experiments to retain expression of the native protein, whereas homozygous Rosa26-tdTomato and Rosa26- DTA mice were used to enhance recombination efficiency.

### Animal procedures

A stock solution of Tamoxifen at 20 mg/mL (Chemodex, T0200) was prepared in a solution of 10% absolute ethanol and 90% olive oil. A dose of 5 mg per 30 g body weight was administered via gavage one day prior to *A. fumigatus* inoculation. TAM citrate- infused chow (400 mg/Kg; Inotiv TD130860) was subsequently provided to the mice for the duration of the experiment. For inoculation of conidia, 0.04 mL of a saline suspension containing 3 x 10^7^ *A. fumigatus* conidia (unless otherwise stated) was administered by the oropharyngeal route, as previously described^63^. Control mice were inoculated with saline. To induce invasive aspergillosis, mice were immunosuppressed by subcutaneous injection of a single dose of triamcinolone acetonide (40 mg/kg body weight)^64^. Survival was monitored for 14 days until mice met pre-specified IACUC-approved early removal criteria.

All mouse studies were conducted in accordance with the Guide for the Care and Use of Laboratory Animals of the National Research Council. The animal use protocol was approved by the Institutional Animal Care and Use Committee (IACUC) at the University of Cincinnati.

### Histology and immunohistochemistry

Harvested lungs were inflated with 4% paraformaldehyde (PFA) through the trachea using gravitational flow, fixed in PFA for 1 h at room temperature, and immersed in 30% sucrose (w/v) overnight at 4°C. The lungs were then embedded in OCT (Fisher, 23730571) and stored at -80°C prior to cryosectioning. For immunohistochemistry, cryosections of 10 μm were soaked in blocking solution (PBS containing 0.1% TritonX- 100, 1.5% goat serum, and 1% bovine serum albumin) for 30 min, followed by overnight incubation at 4⁰C with the primary antibody diluted 1:100 in blocking solution. The primary antibodies used in this study were Myh11(Abcam, AB224804), EpCAM/CD326 (ThermoFisher, 14579185), CD31 (BD Biosciences, 553370), and CD45 (BD Biosciences, 550539). After incubation, sections were washed with PBS and incubated in blocking solution containing a 1:500 dilution of the secondary antibody for 1h at room temperature. The secondary antibodies used were Alexa Fluor 488-conjugated goat anti-rat (ThermoFisher, A11006) or anti-rabbit (ThermoFisher, A11008), and Alexa Fluor 647 goat anti-rat (ThermoFisher, A21247) or anti-rabbit (ThermoFisher, A21244). Sections were then washed, stained with DAPI (Sigma, D9542) for 5 min to label nuclei, and mounted on slides in mounting media (Vector Laboratories, H1700). Epifluorescence images were captured using an Olympus BX43 microscope equipped with an Olympus DP80 camera and connected to an X-Cite 120 fluorescence lamp illuminator. Confocal images were taken with a Leica Stellaris 8 Confocal Microscope. Brightness and contrast adjustments were made using FIJI for epifluorescence images or LAS X software for confocal images. Quantification studies were performed by counting the indicated number of cells in a full longitudinal section of the right lung lobe. At least three biological replicates were quantified per replicate, using two sections separated by 200 µm.

For histopathology, lungs were inflated with PFA, fixed, processed, sectioned, and imaged as described elsewhere^65^. The sections were stained with Gomori’s methenamine silver (GMS), hematoxylin and eosin (HE), or Masson’s trichrome.

### FACS analysis and sorting

Harvested lungs were rinsed in cold PBS and placed in a C-tube (Miltenyi Biotec, 130093237) containing 5 mL of digestion buffer consisting of Dulbecco’s PBS with 0.9mM CaCl2, 600 U/mL Collagenase IV (Worthington Biochemistry, LS004189), 1.2 U/mL Dispase II (Gibco, 17105041) and 30 U/mL DNase I (Sigma, D4527-10KU). The samples were incubated in a water bath at 37°C for 35 min and processed using a MACS™ Dissociator (Miltenyi Biotec, 130-093-235). The homogenate was filtered through a 40 µm cell strainer. Blood cells were eliminated by incubating the samples with Red Cell Lysis Buffer (eBiosciences, 00430054) following the manufacturer’s instructions. The remaining cells were resuspended in FACS buffer (Dulbecco’s PBS with 2x Fetal Bovine Serum) and stained with the following antibodies conjugated with APC (1:100 dilution) for 30 min at room temperature: CD45 (BioLegend, 103112), CD31 (BioLegend, 102409) and EpCAM/CD326 (BioLegend, 118214). Cells were washed with FACS buffer, followed by staining with the viability dye Sytox Blue (Invitrogen, S34857) or Sytox Green (Invitrogen, S34860) at a 1:500 dilution. Flow cytometry analysis was conducted using a BD LSRFortessa and sorting performed on a BD FACSAria II Fusion. Both instruments operated using the FACSDiva software, and analyses were conducted using FlowJo V.10.10. Details of the gating strategies can be found in Fig. S4 and S6.

### RNA sequencing and analysis

For bulk RNAseq analysis, Postn^Lin^ or Pdgfrα-GFP cells were sorted, and the RNA isolated using an RNeasy Plus Micro Kit (Qiagen, 74034) according to the manufacturer’s instructions. RNA integrity was confirmed using the RNA 6000 Pico Kit on an Agilent 2100 Bioanalyzer before sequencing. The cDNA was amplified with the Ovation RNA-Seq System v2 (Tecan Genomics), and libraries were prepared using the Nextera XT DNA Library Preparation Kit (Illumina). Sequencing was performed on a NovaSeq 6000 SP v1.5 (200 cycles), yielding 100 bp paired-end reads with a sequencing depth of 30 million reads per sample. Fastq files from each sample were merged, and quality control of the sequencing reads was conducted. Adapters sequences were trimmed before further analysis. Gene expression analysis was carried out using the *Mus musculus* reference genome GRCm39 (mm39) in CLC Genomics Workbench (Qiagen). Differential expression was assessed between Postn^Lin^ cells treated with *A. fumigatus* and Pdgfrα- GFP cells treated with saline. Expression values were reported as total counts or Reads Per Kilobase per Million mapped reads (RPKM), with a statistical cutoff of a Bonferroni-adjusted P-value ≤ 0.05. Gene ontology (GO) enrichment analysis was performed using the NIH DAVID Bioinformatics Functional Annotation Tool (https://david.ncifcrf.gov) applying a statistical threshold of FDR P-value ≤ 0.01 and ≥ 5 total genes per GO term. GO enrichment and volcano plots were created using RStudio (version 2024.04.2+764) on macOS.

For single-cell RNAseq analysis, sorted pulmonary stromal cells (negative for CD45, CD31 and EpCAM) were collected in a tube containing DMEM (Corning, 10013CV) with 10% FBS at a concentration of 100 cells/μL. The single-cell RNA-Seq assay was performed according to the manufacturer’s instructions (Chromium Next GEM Single Cell 3ʹ Reagent Kits v3.1 (Dual Index), 10x Genomics). Briefly, cells were resuspended in the master mix and loaded together with partitioning oil and gel beads into the chip to generate a gel bead-in-emulsion (GEM). The poly-A RNA from the cell lysate contained in every GEM was reverse transcribed into cDNA, adding an Illumina TruSeq R1 primer sequence, Unique Molecular Identifier (UMI) and the 10x Barcode. The cell barcoded molecules were then cleaned up with Silane DynaBeads and amplified using 14 PCR cycles. Next, full-length, barcoded cDNA was then enzymatically fragmented, sized-selected, adapter-ligated, and amplified for library construction. During the library construction, P5, P7, i7 and i5 sample indexes, and TruSeq Read 2 were added. Samples were pooled and run on the NovaSeq 6000 sequencer with a S4 flow cell using the following sequencing parameters: R1: 28 cycles, i7: 10 cycles, i5: 10 cycles, R2: 90 cycles.

The 10x scRNA-seq data were preprocessed using Cell Ranger software (7.0.0). We used the ‘mkfastq’, ‘count’ and ‘aggr’ commands to process the 10x scRNA-seq output into one cell by gene expression count matrix using default parameters. scRNA-seq data analysis was performed with the Scanpy (1.9.0) package in Python (https://genomebiology.biomedcentral.com/articles/10.1186/s13059-017-1382-0). Genes expressed in fewer than three cells were removed from further analysis. Cells expressing less than 100 and more than 7,000 genes were also removed from further analysis. In addition, cells with a high (≥ 10%) mitochondrial genome transcript ratio were removed. For downstream analysis, we used count per million (CPM) normalization to control for library size differences in cells and transformed those into log (CPM + 1) values. After normalization, the data were then *z*-score normalized for each gene across all cells. We then used the following commands, tl.pca’, ‘pp.neighbors’ and ‘tl.leiden, in Scanpy to partition the single cells into clusters. Differential expression analysis was done using the Wilcoxon rank sum test at the single cell level.

### Data availability

The data sets generated were deposited in the Gene Expression Omnibus (GEO)^66^ with the accession number GSE284270 and GSE284713.

### Statistical analysis

Statistical data analysis was performed using GraphPad Prism v10.3.1.

## Acknowledgements

This work was supported by National Institutes of Health grant R01 R01AI159078 (DSA and OK), R01-HL148598 (OK), and University of Cincinnati Department of Pathology & Laboratory Medicine Pilot Grant to JPGA. SMS were supported by an NIH Training grant T32HL125204 (PIs: Molkentin and Kranias). OK was supported by an American Heart Association Career Development Award CDA34110117. We would like to acknowledge the assistance of the Research Flow Cytometry Facility in the Division of Rheumatology at Cincinnati Children’s Hospital Medical Center.

---

**Figure S1. Dynamics of conidial clearance from the lung.** (A) Kinetics of fungal clearance from the lungs of immunocompetent mice following inoculation with the specified amount of *A. fumigatus* conidia. Values represent the means ± SD from 3 biological replicates. (B) Histological sections of control lungs stained with H&E, Masson’s trichrome and GMS. (C) Paraffin-embedded lung sections from mice inoculated with 3 x 10^7^ *A. fumigatus* conidia. Lungs were harvested at the indicated times, and tissue sections were stained with GMS to highlight fungal elements. Arrows highlight areas containing *A. fumigatus* germlings.

**Figure S2. The appearance of *A. fumigatus*-induced Postn^Lin^ cells is tightly regulated by the TAM-inducible system and is independent of the fungal strain.** Representative fluorescent images show tdTomato+ cells in red and DAP-stained nuclei in blue. (A) Top 3 panels represent control mice, whereas the lower 3 panels represent *A. fumigatus*-inoculated mice. Top left panel: untreated reference control (mice lacking the Postn-MerCreMer and Rosa26-tdTomato insertions); Top middle panel: Postn^Lin^- tracing mice with TAM treatment in the absence of saline mock inoculation; Top right panel: Postn^Lin^-tracing mice mock-inoculated with saline and treated with TAM. Lower left panel: Mice inoculated with 3 x 10^7^ CEA10 conidia without TAM treatment; Lower middle panel: mice inoculated with CEA10 conidia with TAM treatment; Lower right panel: mice inoculated with the same number of heat-killed (HK) CEA10 conidia with TAM treatment. (B) Mice inoculated with the same number of live conidia from the AF293 or H237 clinical isolates of *A. fumigatus*.

**Figure S3. Dose-response analysis to determine the optimal dose of conidia required to elicit fibroblast activation.** Representative histological sections show tdTomato+ cells in the lungs of Postn^Lin^ mice harvested 7 days after OP inoculation with the indicated amounts of *A. fumigatus* conidia. Tissue autofluorescence was enhanced to provide background tissue visualization.

**Figure S4. Postn^Lin^ activated fibroblasts exhibit distinct morphological features.** Representative fluorescent micrographs showing: (A) low-power and (B-D) high-power images of non-activated resident Pdgfrα^Lin^ pulmonary fibroblasts. Arrows on panels B, C, and D highlight Pdgfrα^Lin^ cells (red) surrounding a vessel (V), bronchiole (Br), and alveoli (Al), respectively; (E) low-power and (F-H) high-power images of alveolar regions containing Postn^Lin^ cells embedded within inflammatory foci 7 days after *A. fumigatus* inoculation.

**Figure S5. Cell sorting strategy used for quantification of coexpression markers.** (A-D) Representative flow cytometry gating strategy to differentiate Postn^Lin^ cells (tdTomato+) from non-stromal cells (APC+): (A) Cell gating, (B & C) singlet identification, and (D) selection of live cells. The resulting tdTomato+ population (rightward scatter) was analyzed for expression of (E) EpCAM, (F) CD31, and (G) CD45, (upward scatter), labeled with allophycocyanin (APC). The percentage of tdTomato+ cells that are positive for these markers are indicated in the upper right quadrant. FSC: Forward-scatter, SCC: Side-scatter, A: Area, H: Height, W: Width.

**Figure S6. Cell sorting strategy for isolating tdTomato+ stromal cells for bulk RNA-seq.** (A) Cell gating, (B-C) singlet identification, and (D) live cell selection. (E) The resulting non-stromal population was excluded by removing CD31+ (endothelial), EpCAM+ (epithelial), and CD45+ (hematopoietic) cells labeled with allophycocyanin (APC). (F) The remaining tdTomato+ stromal population (rightward scatter) was sorted for RNA isolation and bulk RNA-seq. FSC: Forward-scatter, SCC: Side-scatter, A: Area, H: Height, W: Width.

**Table S1. List of deferentially expressed genes from bulk-RNAseq analysis**

**Table S2. List of GO terms for Upregulated genes.**

**Table S3. List of GO terms for Downregulated genes.**

**Table S4. Functional classification of DEGs in *A. fumigatus* induced clusters Activated^Alv^ and Activated^Pbr^.**

