## Supplementary figures and images for "Pulmonary fibroblast activation during *Aspergillus fumigatus* infection enhances lung defense via immunomodulation and tissue remodeling"

### Supplemental Figures 1-6

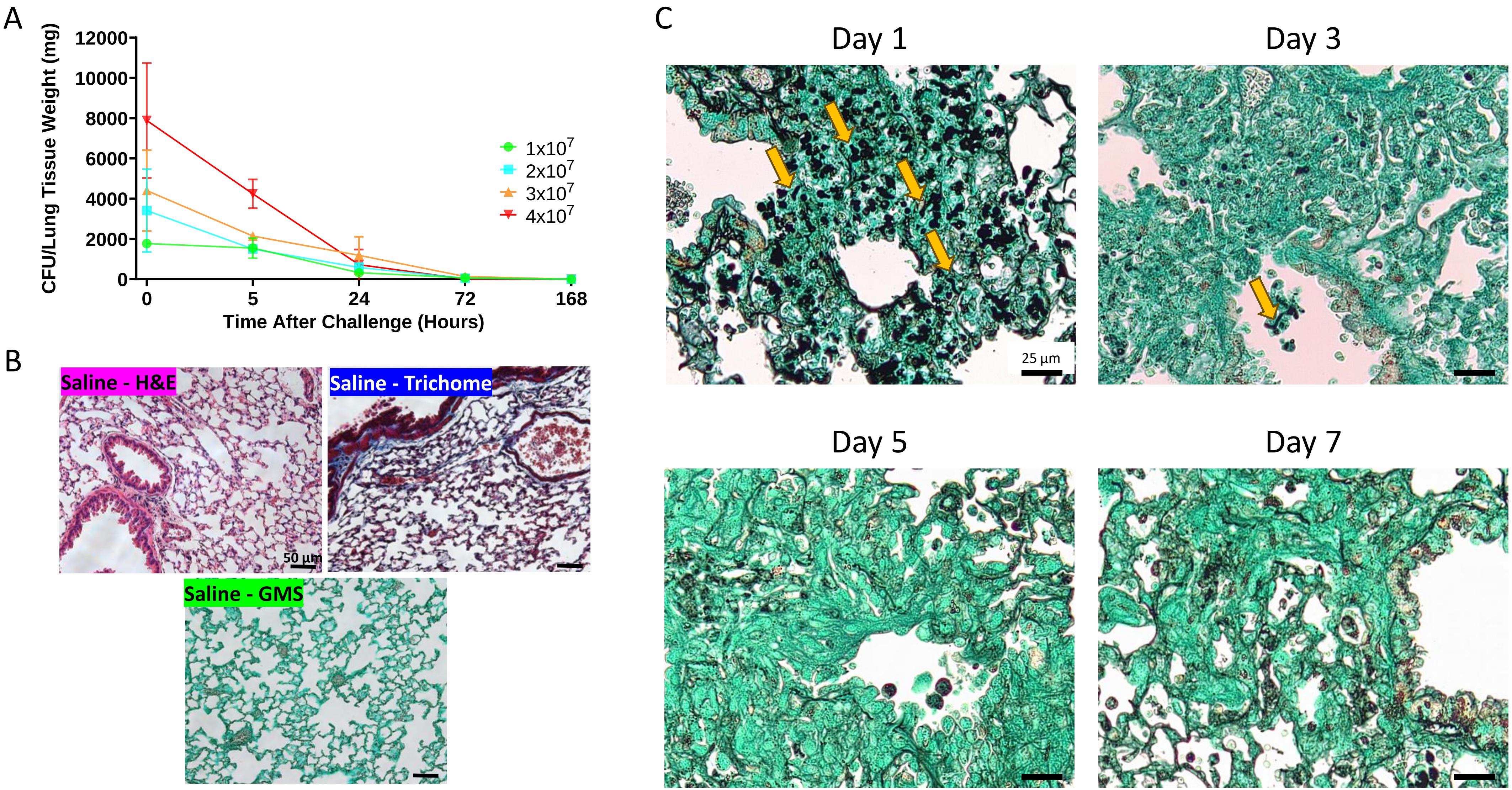

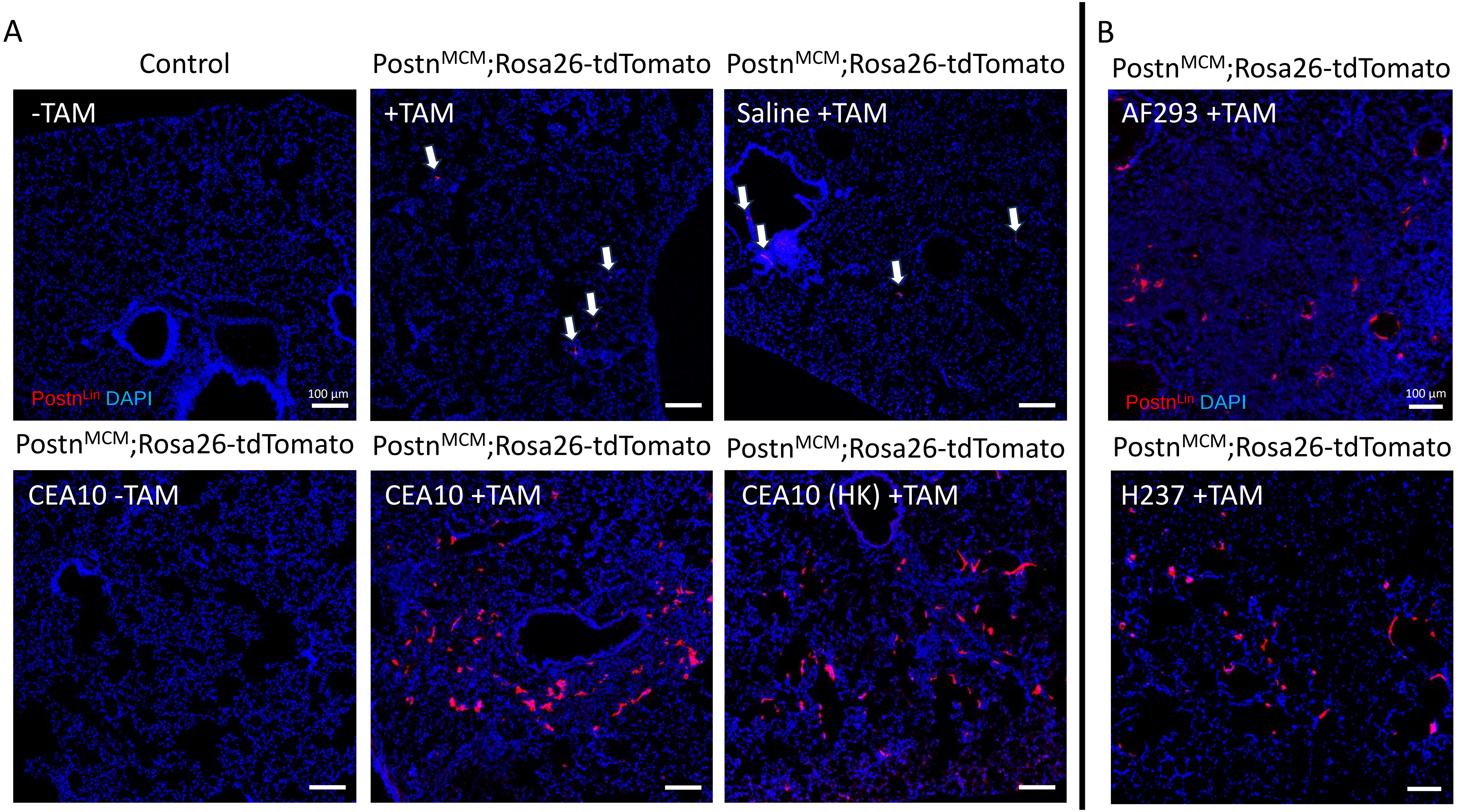

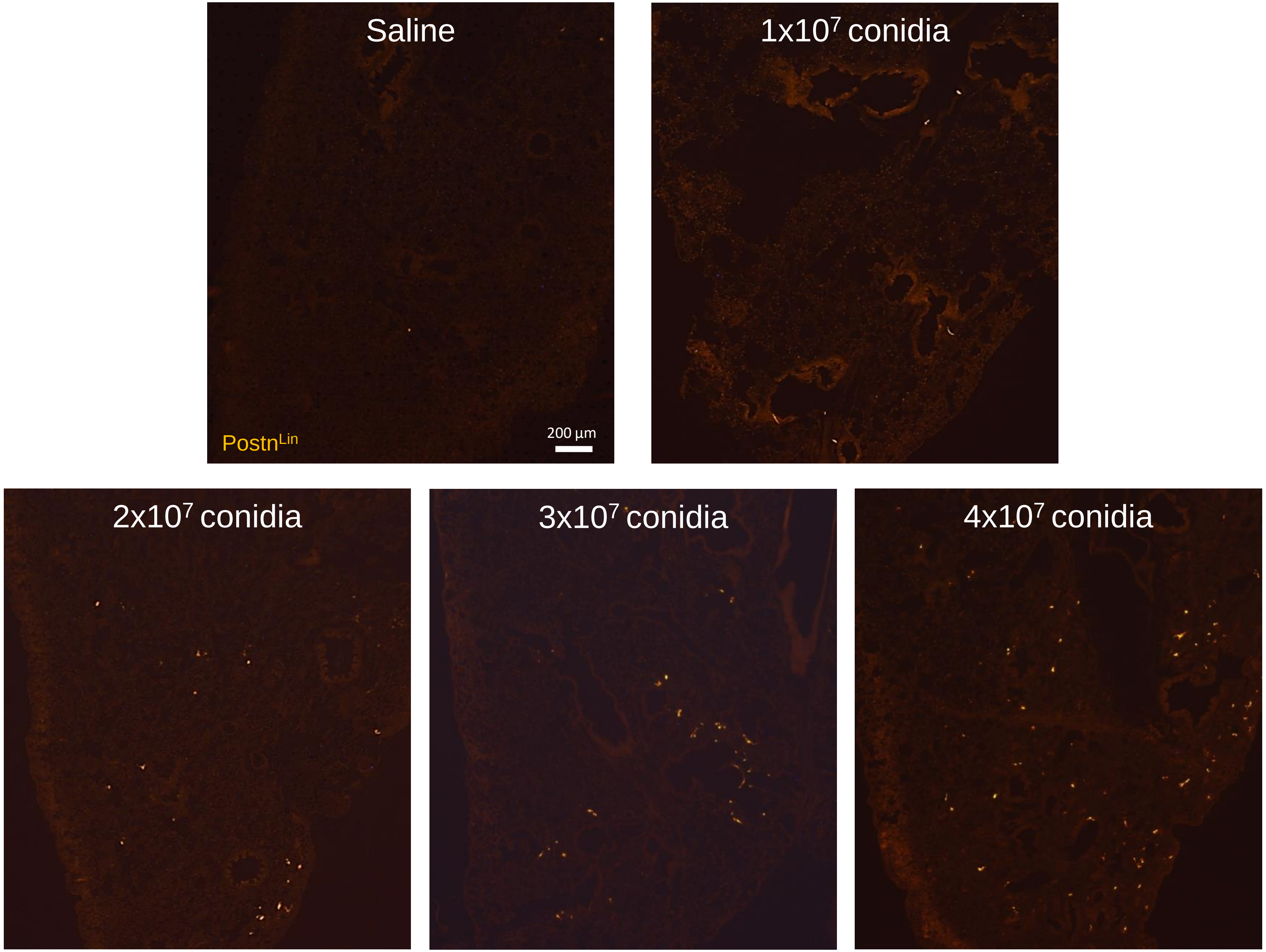

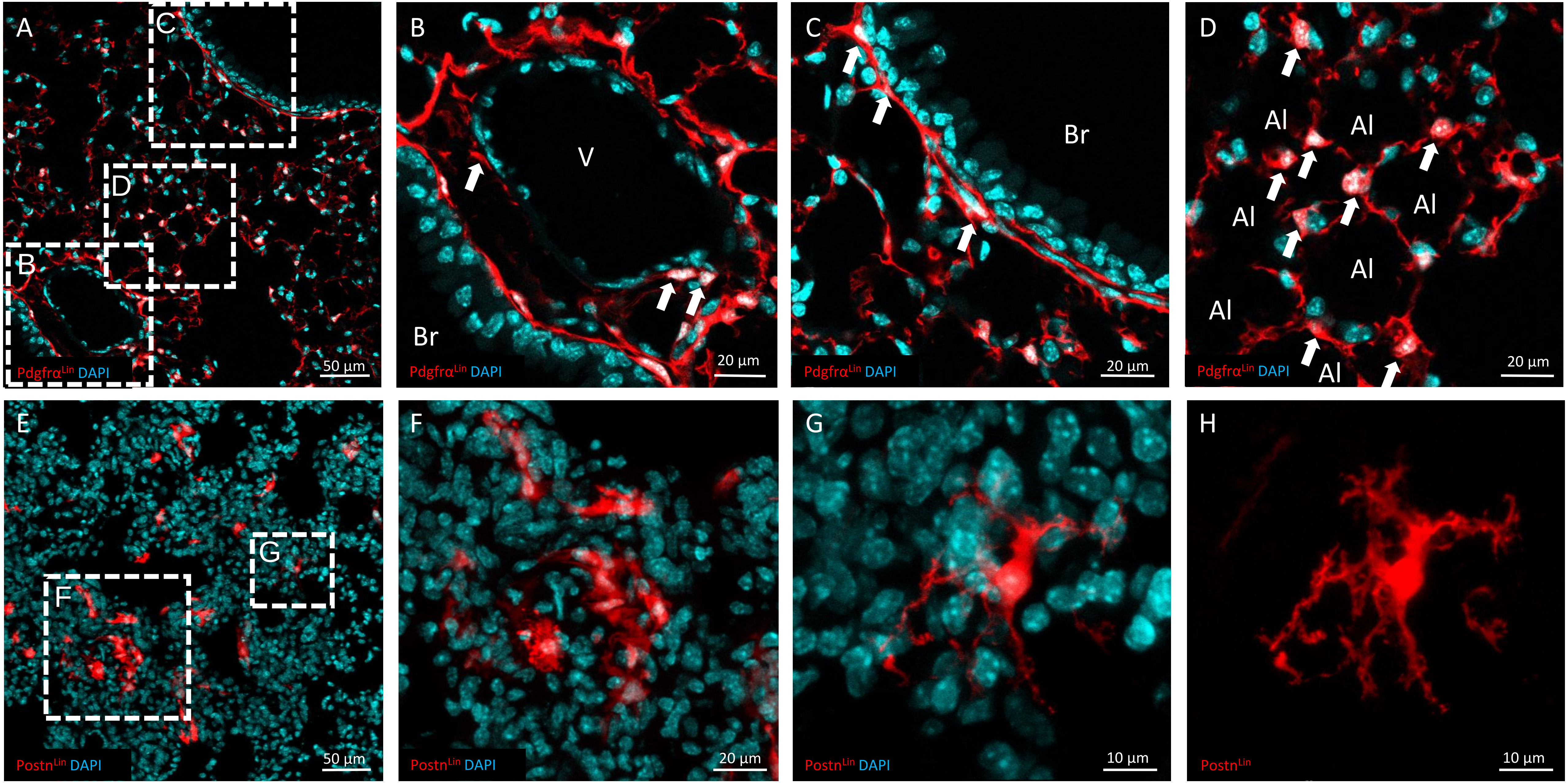

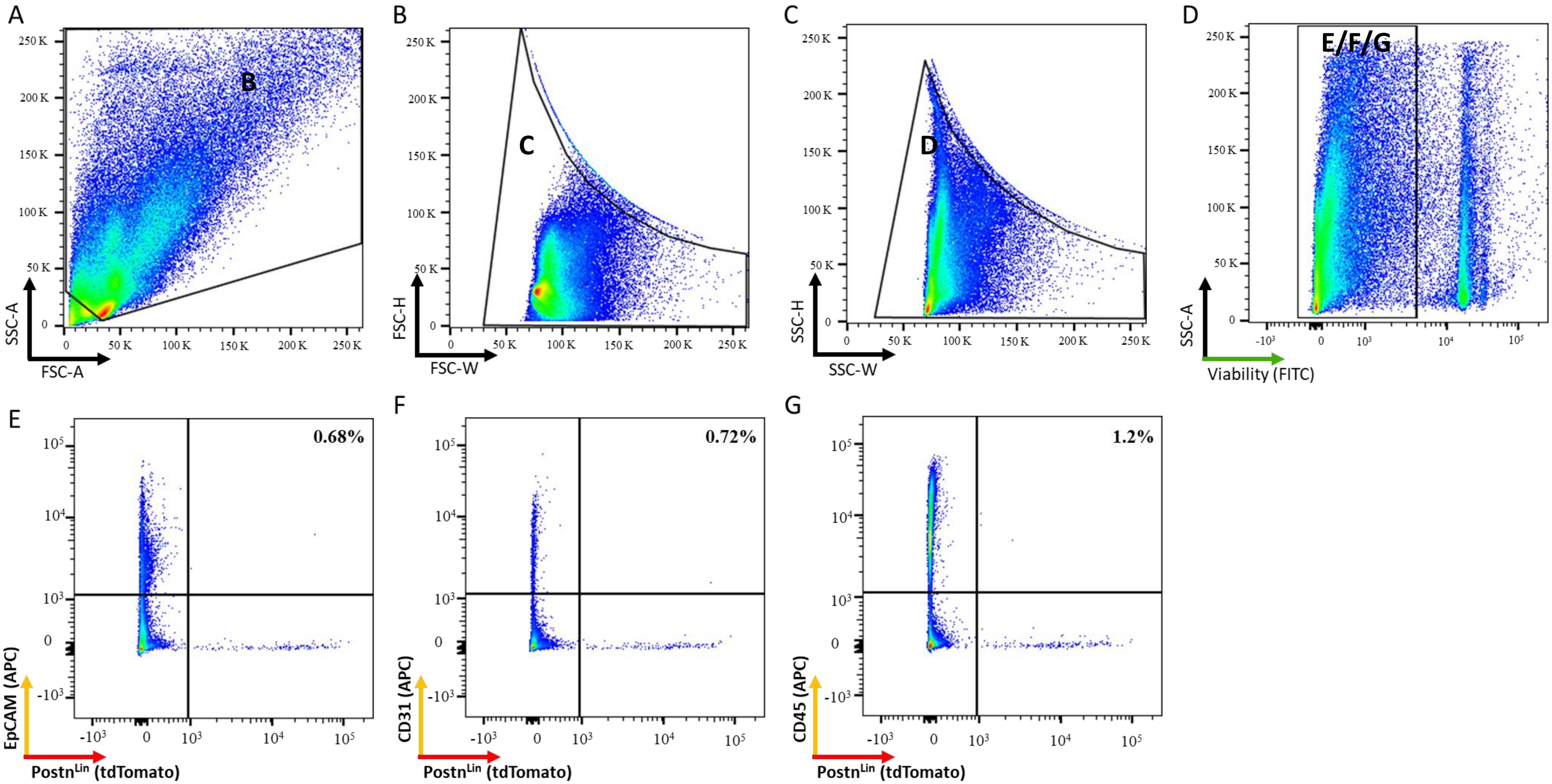

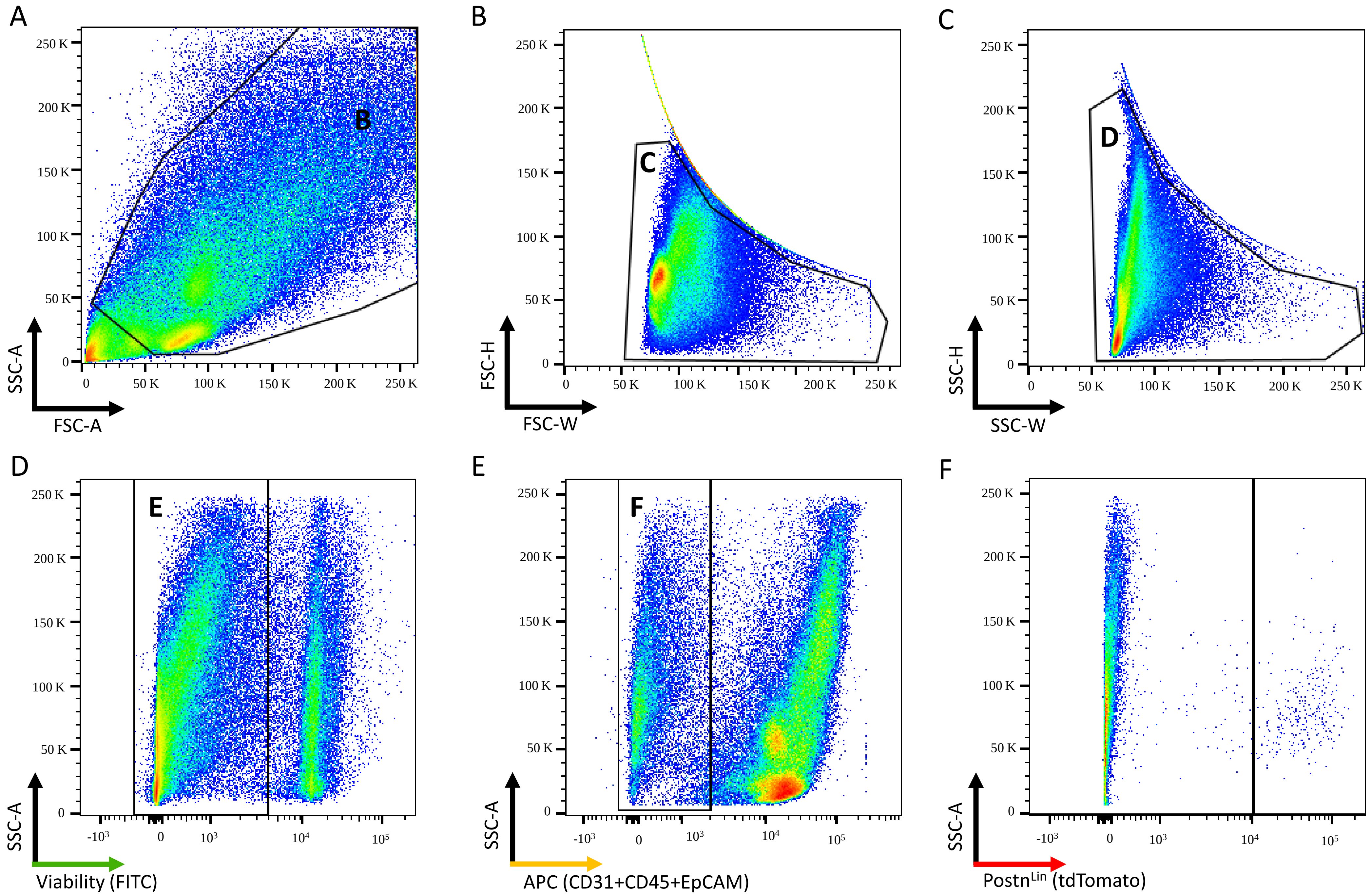
